# Hem25p is a mitochondrial IPP transporter

**DOI:** 10.1101/2023.03.14.532620

**Authors:** Jonathan Tai, Rachel M. Guerra, Sean W. Rogers, Zixiang Fang, Laura K. Muehlbauer, Evgenia Shishkova, Katherine A. Overmyer, Joshua J. Coon, David J. Pagliarini

## Abstract

Coenzyme Q (CoQ, ubiquinone) is an essential cellular cofactor comprised of a redox-active quinone head group and a long hydrophobic polyisoprene tail. How mitochondria access cytosolic isoprenoids for CoQ biosynthesis is a longstanding mystery. Here, via a combination of genetic screening, metabolic tracing, and targeted uptake assays, we reveal that Hem25p—a mitochondrial glycine transporter required for heme biosynthesis—doubles as an isopentenyl pyrophosphate (IPP) transporter in *Saccharomyces cerevisiae*. Mitochondria lacking Hem25p fail to efficiently incorporate IPP into early CoQ precursors, leading to loss of CoQ and turnover of CoQ biosynthetic proteins. Expression of Hem25p in *Escherichia coli* enables robust IPP uptake demonstrating that Hem25p is sufficient for IPP transport. Collectively, our work reveals that Hem25p drives the bulk of mitochondrial isoprenoid transport for CoQ biosynthesis in yeast.

## Introduction

Coenzyme Q (CoQ, ubiquinone) is a redox-active lipid that functions as an essential cofactor for multiple cellular processes, including oxidative phosphorylation, pyrimidine biosynthesis, fatty acid oxidation, and ferroptotic defense^1–4^. Deficiencies in CoQ underlie multiple human pathologies, all of which have limited therapeutic options^5,6^. Despite being discovered 65 years ago, significant gaps in knowledge persist for how CoQ is synthesized and distributed throughout the cell, thereby limiting the development of interventional strategies^7–9^.

CoQ is comprised of a fully-substituted quinone ring and a long hydrophobic tail consisting of repeating isoprene units (Fig. 1a). CoQ is synthesized on the matrix side of the inner mitochondrial membrane beginning with the precursors 4-hydroxybenzoate (4-HB) and isoprenoid pyrophosphates^10,11^. These precursors are each formed in the cytosol: 4-HB is derived from tyrosine or the shikimate pathway, and the isoprenoid pyrophosphates are produced by the mevalonate pathway ^8^. Prior work has shown that mitochondria are able to import 4-HB and the isoprenoid isopentenyl pyrophosphate (IPP) for CoQ biosynthesis *in vitro*^12,13^; however, the protein(s) that enable this transport have remained elusive for decades. Previous efforts to identify proteins necessary for CoQ production—largely by leveraging the inability of yeast lacking CoQ to grow in respiratory conditions—have failed to identify candidate transporters^8,14,15^, suggesting that there is either redundancy in the system or that the requisite transporter is essential for growth under common conditions^16^.

**Fig. 1.**
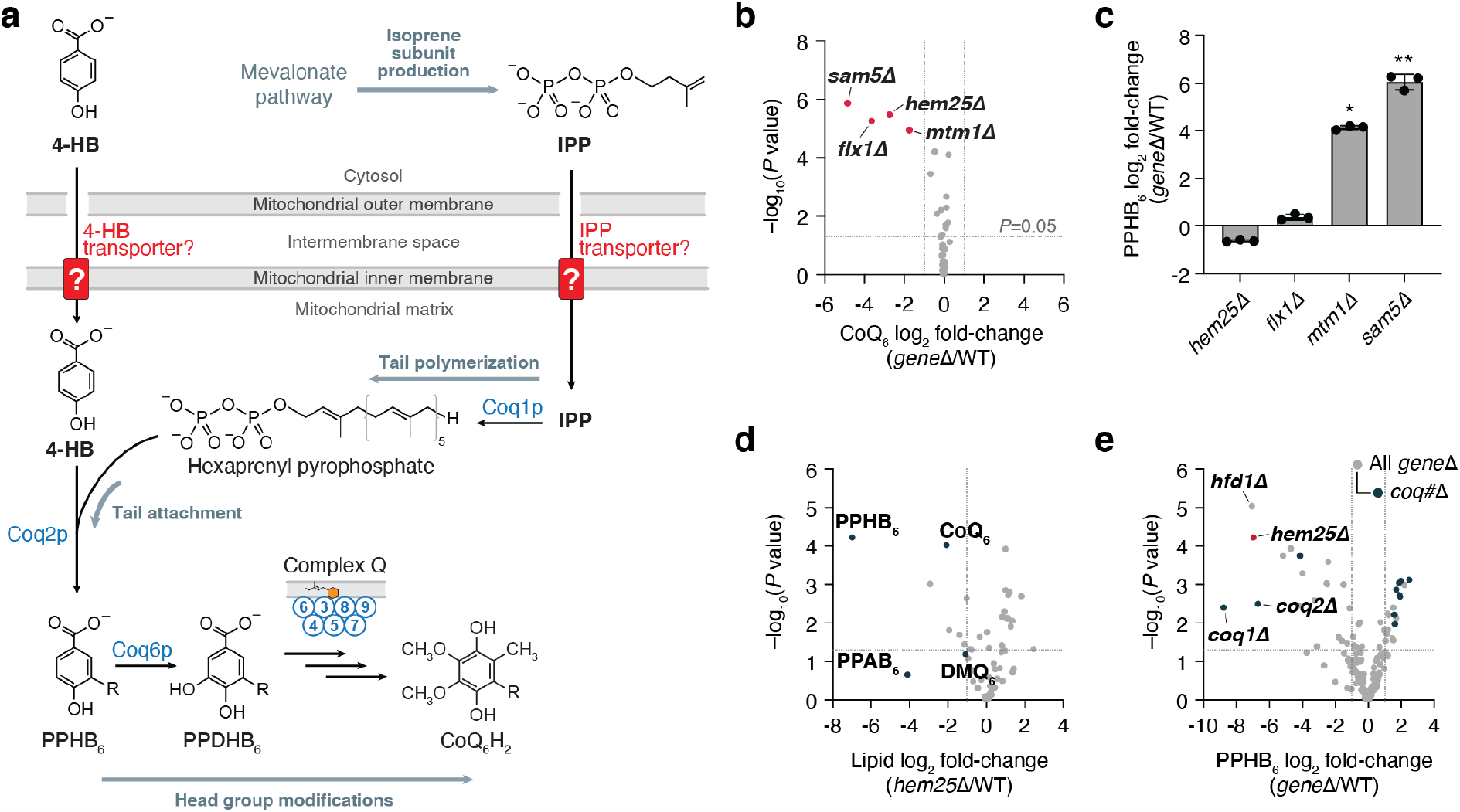
A targeted genetic screen identifies Hem25p as a potential transporter of CoQ precursors. **a**, Schematic of CoQ biosynthesis in *S. cerevisiae*. 4-HB, 4-hydroxybenzoate; IPP, isopentenyl pyrophosphate; PPHB_6_, polyprenyl-hydroxybenzoate; PPDHB6, polyprenyl-dihydroxybenzoate. **b**, Relative CoQ abundance in all *gene*Δ strains compared to WT verses statistical significance. Hits with significantly decreased levels of CoQ are highlighted (mean, *n*=3 independent samples, two-sides Student’s *t-*test). **c**, Relative PPHB levels in each of the hits from (**b**) (^*^ *p=*1.02 × 10^−5^ WT vs *mtm1*Δ, ^* *^ *p=*0.0012 WT vs *sam5*Δ; mean ± SD, *n*=3 independent samples). **d**, Relative lipid abundances in *hem25*Δ yeast compared to WT verses statistical significance with CoQ and CoQ biosynthetic intermediates highlighted. PPAB6, polyprenyl-aminobenzoate; DMQ6, demethoxy-coenzyme Q. **e**, Relative PPHB abundances verses statistical significance across all Y3K *gene*Δ strains with *hem25*Δ, *hfd1*Δ, and coq#Δ strains highlighted. For panels (**d**) and (**e**), raw data from the Y3K dataset^30^ (respiration-RDR condition) is displayed as the mean from 3 independent samples with two-sided Student’s *t-*test used for both panels.

## Results

### A targeted screen for transporters in CoQ biosynthesis identifies Hem25p

To identify mitochondrial transporters important for CoQ production, we created a custom panel of *S. cerevisiae* gene knockout (*gene*Δ) strains each lacking an established or potential mitochondrial transporter. This panel comprised *gene*Δ strains for all 35 members of the SLC25 family of mitochondrial transporters, which are responsible for nearly all known metabolite transport into mitochondria^17^, and seven poorly characterized mitochondrial inner membrane proteins that might possess transporter function^18,19^. We grew strains in triplicate under fermentative conditions and monitored their ability to produce CoQ via targeted electrochemical detection coupled to HPLC (HPLC-ECD). Using this strategy, we identified four strains with significantly decreased CoQ levels (Fig. 1b). Of these strains, two lacked transporters (Flx1p and Sam5p) known to carry substrates required for the later, head-group modifying steps of CoQ biosynthesis (the cofactors FAD and S-adenosylmethionine, respectively)^20–23^. The third lacked Mtm1p, an uncharacterized mitochondrial transporter. Cells lacking Mtm1p demonstrate elevated mitochondrial manganese levels and disrupted iron-sulfur cluster biogenesis, both of which can compromise late CoQ biosynthesis^7,24–26^. The final strain lacked Hem25p, a glycine carrier with no established connection to CoQ^27,28^.

To further prioritize these candidates, we then measured the levels of polyprenyl-hydroxybenzoate (PPHB). PPHB and, to a lesser extent polyprenyl-aminobenzoate (PPAB), are the first intermediates of CoQ produced within mitochondria and are known to accumulate when the downstream pathway is disrupted. We reasoned that loss of cofactor transporters important for enzymes in the later stages of CoQ biosynthesis would likewise cause an accumulation of PPHB, whereas loss of a 4-HB or IPP transporter would prevent PPHB formation^29^. Of the four candidates, only the *hem25*Δ strain had decreased levels of PPHB (Fig. 1c). This finding is consistent with data from our recent systematic analyses of mitochondrial protein functions, in which the *hem25*Δ strain exhibited decreased levels of CoQ and the prenylated CoQ precursors PPHB and PPAB (Fig. 1d)^30^. In reanalyzing data from all 176 strains in that study (which included few transporter KO strains), *hem25*Δ exhibited PPHB depletion comparable to strains lacking the established early-stage CoQ proteins Hfd1p (which produces 4-HB) and Coq1p and Coq2p (which collectively produce PPHB and PPAB) (Fig. 1e). Intriguingly, large-scale chemical genomic screens have also linked *HEM25* to the mevalonate pathway and statin sensitivity (Supplementary Fig. 1a,b), suggesting unexplored connections to isoprenoid biology^31^. Thus, we proceeded to investigate Hem25p in biochemical depth.

### Hem25p has distinct roles in heme and CoQ production

Hem25p was previously established as a mitochondrial glycine transporter necessary for heme biosynthesis (Fig. 2a)^27,28^. Imported glycine condenses with succinyl-CoA to form the heme precursor aminolevulinate (ALA). ALA is then exported out of the mitochondrial matrix to the cytosol where it proceeds along the heme biosynthetic pathway. Yeast lacking Hem25p can still produce low levels of heme^28^, suggesting that mitochondria contain a secondary glycine carrier. However, its identity remains unknown.

**Fig. 2.**
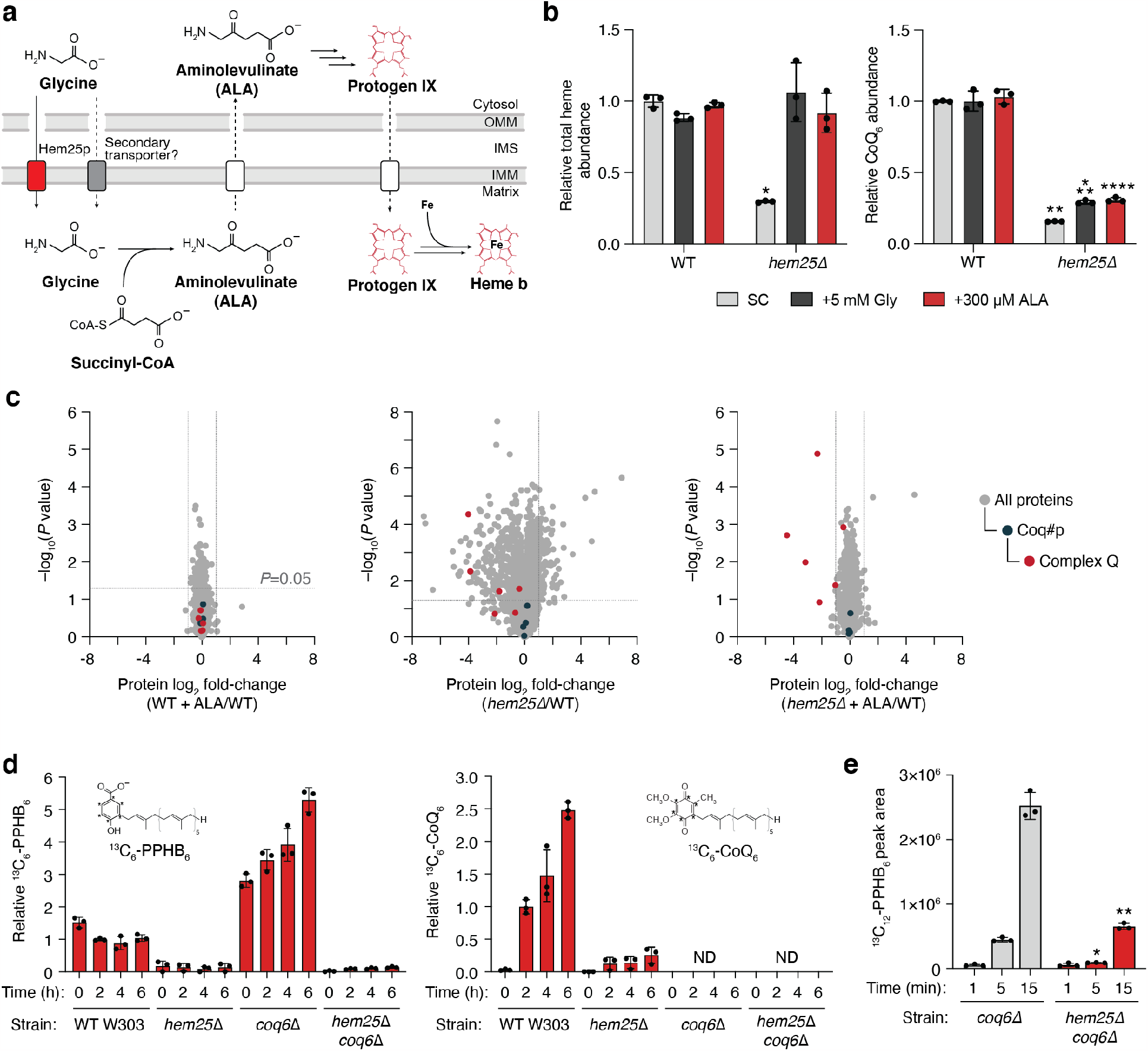
Hem25p drives CoQ production independently of its role in heme biosynthesis. **a**, Overview of heme biosynthesis and the role of Hem25p. **b**, Relative total heme and CoQ abundances in WT and *hem25*Δ yeast grown in synthetic complete (SC) media with and without glycine and aminolevulinate (ALA) supplementation (^*^ *p=*1.01 ’ 10^−5^ WT SC heme vs *hem25*Δ SC heme, ^**^*p=*7.56× 10^−10^ WT SC CoQ vs *hem25*Δ SC CoQ, ^***^*p=*8.47× 10^−8^ WT SC CoQ vs *hem25*Δ +Gly CoQ, ^****^*p*=2.22× 10^−7^ WT SC CoQ vs *hem25*Δ +ALA CoQ; mean ± SD, *n*=3 independent samples). **c**, Relative protein abundances of WT cells with ALA, *hem25*Δ cells without ALA, and *hem25*Δ cells with ALA supplementation compared to WT cells without ALA supplementation verses statistical significance. CoQ biosynthetic proteins and complex Q proteins are highlighted (mean, *n*=3 independent samples, two-sides Student’s *t-*test). **d**, Relative abundance of *de novo* synthesized ^13^C6-PPHB and ^13^C6-CoQ in WT, *hem25*Δ, *coq6*Δ, and *coq6*Δ*hem25*Δ cells grown in -*p*ABA SD, 3% (w/v) glycerol, and 50 *μ*M ^13^C_6_-4-HB (mean ± SD, *n*=3 independent samples); ND, not detected. **e**, Normalized abundance of *de novo* synthesized ^13^C_12_-PPHB in isolated *coq6*Δ and *coq6*Δ*hem25*Δ mitochondria (^*^*p*=5.73× 10^−5^ WT vs *hem25*Δ at 5 min, ^**^*p*=0.0001 WT vs *hem25*Δ at 15 min; mean ± SD, *n*=3 independent samples).

To determine if the effect of Hem25p on CoQ levels is related to its role in heme production, we supplemented the *hem25*Δ strain with either ALA or high levels of glycine, both of which have been shown to bypass the requirement of Hem25p for heme production^27,28^. Without supplementation, *hem25*Δ cells had markedly reduced levels of heme and CoQ (Fig. 2b). Consistent with prior work, both glycine and ALA supplementation fully rescued heme in the *hem25*Δ strain to wild-type (WT) levels. However, supplementation had minimal effect on CoQ levels, demonstrating distinct roles for this transporter. Notably, the residual levels of CoQ in *hem25*Δ suggests that other transporters may enable limited isoprenoid uptake. This may explain why Hem25p has been difficult to identify by screens for respiratory incompetency, as very little CoQ is required for yeast to survive on non-fermentable carbon sources^32^.

To further probe Hem25p’s distinct roles in heme and CoQ biosynthesis, we performed whole-cell proteomic analyses on the *hem25*Δ strain. Unsurprisingly, disruption of Hem25p resulted in widespread protein changes, with 185 proteins showing greater than two-fold decrease in abundance versus WT cells. Of these significantly decreased proteins, 121 were mitochondrial, including select CoQ biosynthetic proteins (Fig. 2c). A similar proteomic response was seen in our previous large-scale proteomics analyses (Supplemtary Fig. 2a)^30^. Remarkably, *hem25*Δ cells supplemented with ALA exhibited a decrease in only four proteins, all of which are enzymes in the later stages of the CoQ biosynthesis pathway (Fig. 2c). Furthermore, this reduction in CoQ biosynthetic proteins was independent of transcript levels (Supplementary Fig. 2b). Previous studies have demonstrated that CoQ or a late CoQ intermediate is essential for stabilizing the CoQ biosynthetic complex (complex Q, CoQ synthome) responsible for the later, head-group modifying reactions and that multiple subunits of this complex are degraded in its absence^33,34^. These results suggest that Hem25p supports the production of CoQ or a stabilizing intermediate, consistent with a role in transporting IPP or 4-HB. Importantly, our proteomic analyses showed an overall decrease in respiratory chain complexes in *hem25*Δ cells that is completely restored to WT levels upon ALA supplementation (Supplementary Fig. 2c). This is consistent with previous reports of Hem25p’s effect on respiratory chain stability, and further demonstrates Hem25p’s distinct roles in heme and CoQ biosynthesis^27,35^.

To assess Hem25p’s contribution to CoQ production, we monitored the incorporation of heavy isotope-labeled 4-HB ([*phenyl*-^13^C]-4-HB) into the CoQ pathway in *hem25*Δ cells supplemented with ALA using a custom mass spectrometry (MS) analysis. Incorporation of labeled 4-HB into PPHB and CoQ was markedly decreased in the *hem25*Δ strain when compared to the WT strain, again suggesting that Hem25p enables PPHB production (Fig. 2d). To minimize any potentially confounding results from decreased complex Q levels in *hem25*Δ, we also compared these results to those from a *coq6*Δ background strain. Coq6p, a hydroxylase, is the first enzyme to act upon PPHB once formed. Yeast lacking Coq6p are unable to hydroxylate PPHB, leading to decreased levels of complex Q proteins, disrupted CoQ biosynthesis, and an accumulation of labeled PPHB^20,34^. This strain thus enables us to isolate a potential role for Hem25p upstream of Coq6p. Indeed, the *coq6*Δ PPHB accumulation was greatly decreased in the *coq6*Δ*hem25*Δ strain, providing additional evidence that Hem25p enables PPHB production, likely by providing one or both precursors required for its formation (Fig. 2d).

To directly measure Hem25p’s contribution to CoQ biosynthesis *in vitro*, we cultured the *coq6*Δ and *coq6*Δ*hem25*Δ strains with ALA, isolated their mitochondria, and monitored their ability to form PPHB in the presence of 4-HB, MgCl_2_, and [1,2-^13^C]-IPP. Consistent with our whole cell results, disruption of Hem25p greatly reduced the ability of mitochondria to generate ^13^C_12_-PPHB (Fig. 2e). Thus, even with ALA supplementation, cells and mitochondria lacking Hem25p exhibit a major deficiency in generating CoQ and CoQ intermediates. Collectively, these results show that Hem25p directly contributes to CoQ biosynthesis and that this role is independent of its established function in glycine transport and heme biosynthesis.

### Hem25p transports isoprenes in bacteria

Our results above suggest a model whereby Hem25p transports a precursor— either 4-HB or IPP—into the mitochondrial matrix for CoQ biosynthesis. To test this directly, we generated a construct encoding an in-frame fusion of the periplasmic maltose-binding protein (MBP) gene with *HEM25*, enabling expression at the *E. coli* plasma membrane^36,37^. Control (empty vector) *E. coli* cells, or those expressing MBP-Yhm2p, the mitochondrial citrate and oxoglutarate carrier^38^, exhibited little to no ability to take up [1-14C]-IPP (Fig. 3a and Supplementary Fig. 3b,c). However, cells expressing MBP-Hem25p demonstrated a clear, time-dependent [1-^14^C]-IPP uptake that was inhibited by excess unlabeled IPP (Fig. 3a). This IPP import was saturatable with a Michaelis constant (KM) of ∼11 *μ*M (Fig. 3c and Supplementary Fig. 3a). In contrast, no 4-HB import was detected in cells expressing MBP-Hem25p (Fig. 3b). Uptake of 4-HB was only seen when the known bacterial 4-HB transporter PcaK was expressed (Supplementary Fig. 3b)^39^.

**Fig 3.**
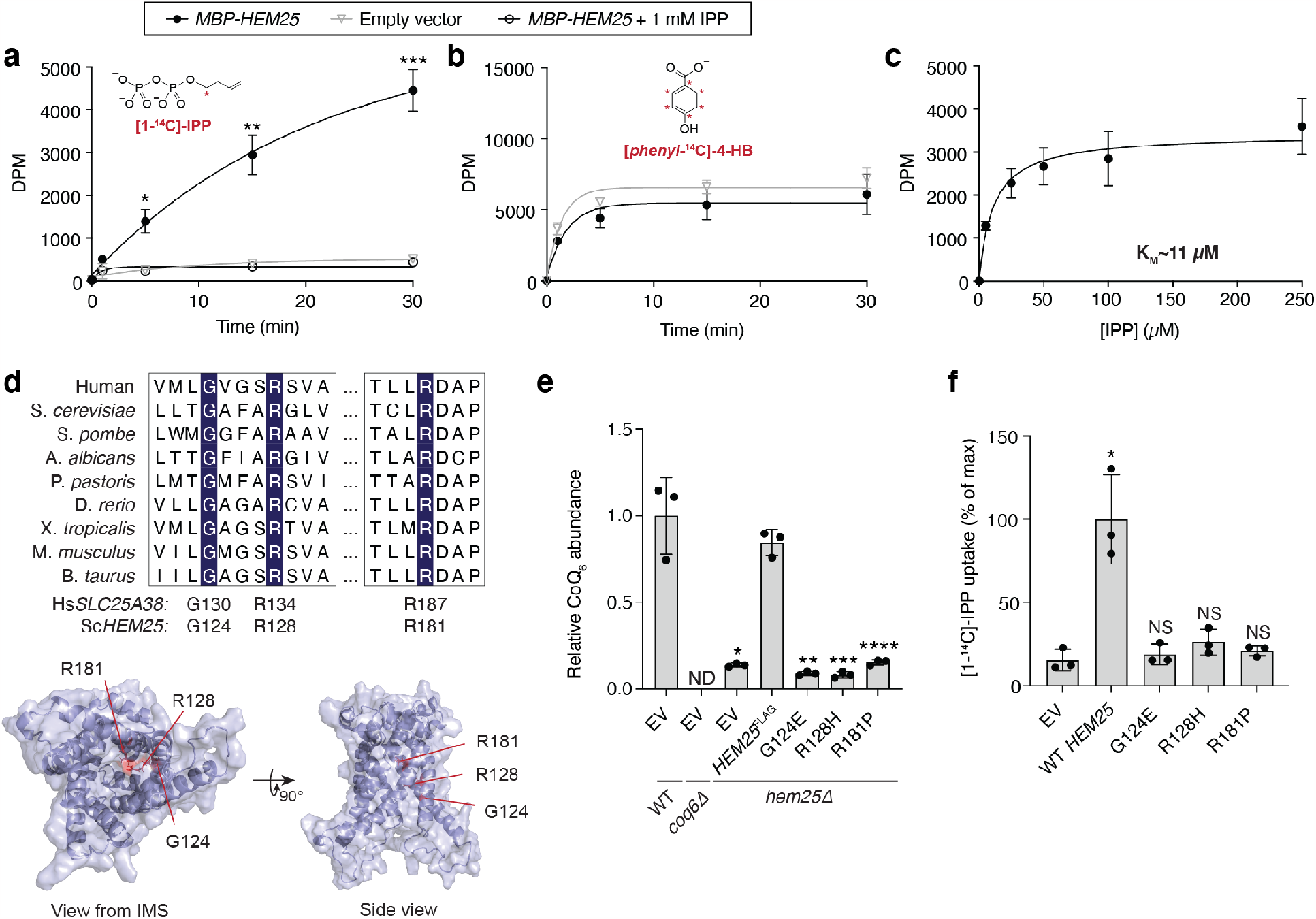
Bacteria expressing Hem25p import IPP. **a**, Time course of 50 *μ*M [1-^14^C]-IPP uptake by *E. coli* cells expressing MBP-Hem25p. Cells carrying the empty expression vector and competition by excess unlabeled IPP were used as controls (^*^*p=*0.0022, ^**^*p=*0.0006, ^***^*p=*0.0001 empty vector vs *MBP-HEM25*, mean ± SD from 3 independent samples); DPM, disintegrations per minute. **b**, Time course of 50 *μ*M [*phenyl*-^14^C]-4-HB uptake by *E. coli* cells expressing MBP-Hem25p. Cells carrying the empty expression were used as controls. Data reflect the mean ± SD from 3 independent samples. **c**, Steady-state kinetics of [1-^14^C]-IPP uptake. For each time point, the corresponding empty vector control was subtracted. Data reflect the mean ± SD from 3 independent samples. **d**, Top, multiple sequence alignment of *HEM25* orthologs with residues mutated in congenital sideroblastic anemia highlighted. Bottom, predicted structure of Hem25p showing the location of the disease-related residues; IMS, intermembrane space. **e**, Relative CoQ abundances in *hem25*Δ yeast carrying WT and mutant *HEM25*^FLAG^ constructs. Levels are relative to WT yeast carrying the empty expression vector (^*^*p=*8.58× 10^−5, **^*p=*6.67×10^−5, ***^*p=*6.78× 10^−5, ****^*p=*9.73× 10^−5^ *HEM25*^FLAG^ vs mutants or empty vector, mean ± SD from 3 independent samples); ND, not detected. **f**, Relative [1-^14^C]-IPP uptake by *E. coli* cells expressing WT and mutant MBP*-*Hem25p. Uptake levels reflect 30 minutes of incubation time and are relative to that of WT MBP-Hem25p (^*^*p=*0.006 empty vector vs WT *HEM25*, mean ± SD, 3 independent samples); NS, not significant.

To ensure that IPP uptake was the result of Hem25p function, we generated three Hem25p point mutants: G124, R128, and R181. All three residues are highly conserved in *HEM25* orthologs, with R128 and R181 located in the common substrate binding site of mitochondrial carriers (Fig. 3d)^40^. When expressed from the endogenous *HEM25* promoter, none of the three mutants was able to rescue CoQ levels (Fig. 3e and Supplementary Fig. 3d,e). Constitutive overexpression of the mutants resulted in only a partial rescue in the R128H and R181P mutants (Supplementary Fig. 3f,g), suggesting that these residues are important for Hem25p’s IPP transport function. Consistent with our CoQ rescue observations, none of the mutants demonstrated IPP transport when expressed in bacteria (Fig. 3f and Supplementary Fig. 3h,i). Taken together, our results show that Hem25p is sufficient to transport IPP into the mitochondrial matrix for CoQ biosynthesis.

### Hem25p’s role in CoQ biosynthesis is restricted to fungi

Hem25p exhibits high overall sequence conservation with human SLC25A38, an established glycine transporter whose disruption results in congenital sideroblastic anemia^41,42^. Given Hem25p’s function in yeast CoQ biosynthesis, we next tested whether the human SLC25A38 also doubles as an IPP transporter. Expression of the human SLC25A38 in the *hem25*Δ strain restored total heme levels, but had no effect on CoQ (Fig. 4a and Supplementary Fig. 4a). Consistently, when expressed in bacteria, MBP-SLC25A38 failed to transport IPP (Fig. 4b). Thus, despite sharing many of the same substrate binding residues as Hem25p, only the glycine transport function is conserved in human SLC25A38. This is further supported by our recent MITOMICS dataset, whereby the human HAP1 cells lacking *SLC25A38* had normal levels of CoQ and complex Q members (Supplementary Fig. 4b,c)^43^.

**Fig. 4.**
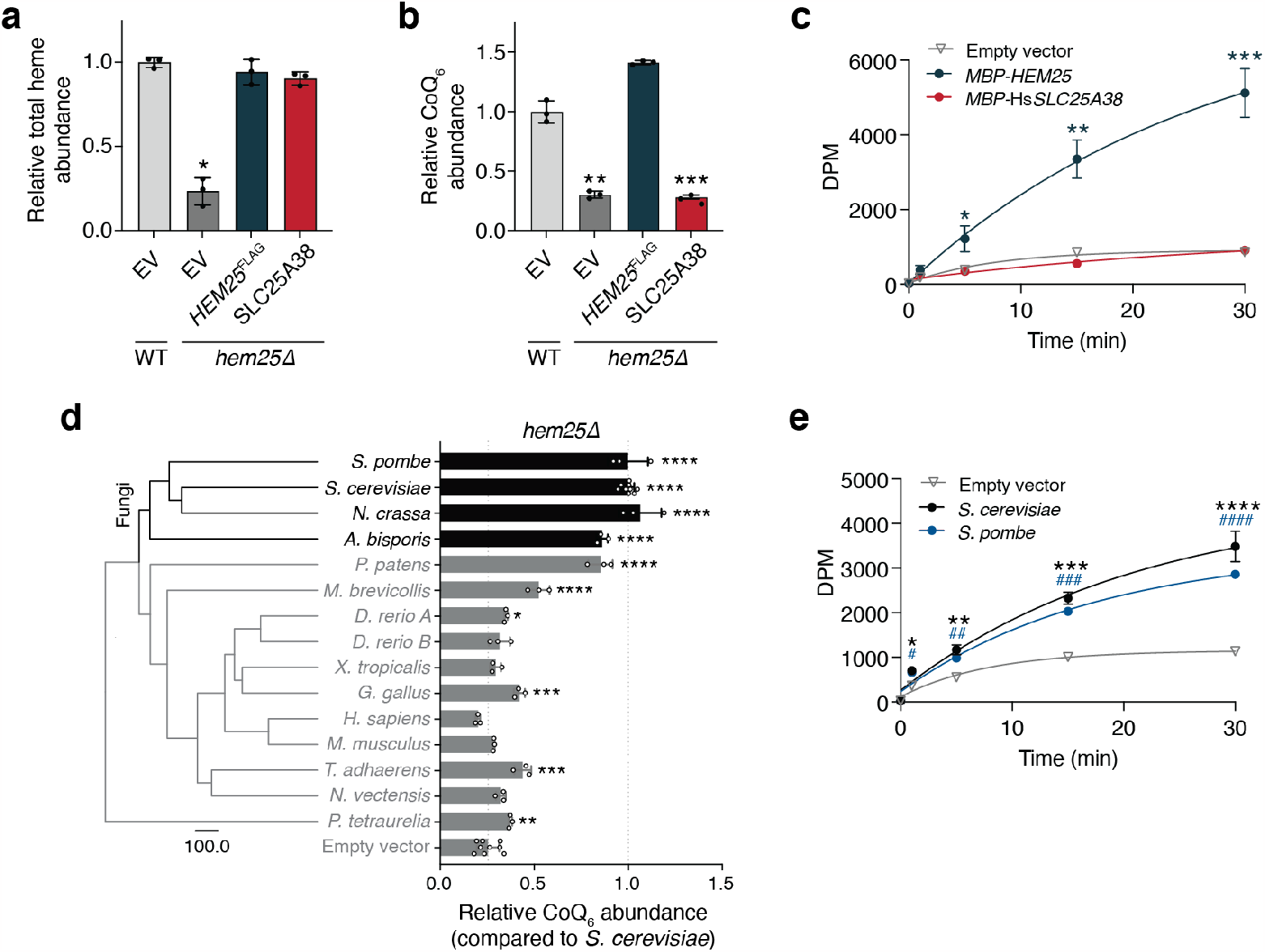
Hem25p’s role in CoQ biosynthesis is conserved in fungi. **a**, Heme and **b**, CoQ rescue by SLC25A38 in *hem25*Δ yeast. Levels are relative to the WT strain carrying the empty expression vector (^*^*p=*0.0001 ^**^*p=*0.0002, ^***^*p=*0.0001, mean ± SD, 3 independent samples). **c**, Time course of 50 *μ*M [1-^14^C]-IPP uptake by *E. coli* cells expressing MBP-SLC25A38 and MBP-Hem25p. Uptake by cells carrying the empty expression vector is included as a control (^*^*p=*0.0147, ^**^*p=*0.0011, ^***^*p=*0.0003 empty vector vs *MBP-HEM25*, mean ± SD, *n=*3 independent samples). **d**, Phylogenetic analysis of Hem25p orthologs, and relative CoQ abundance in *hem25*Δ cells expressing different Hem25p orthologs. Levels are relative to that of the *S. cerevisiae* Hem25p (^****^*p*<0.0001, ^***^*p*<0.001, ^**^*p*<0.01, ^*^*p*<0.05 vs empty vector; mean ± SD, at least 3 independent samples). **e**, Time course of [1-^14^C]-IPP uptake by *E. coli* cells expressing *S. cerevisiae* Hem25p, *S. pombe* Hem25p, or the empty vector. (^*^*p=*0.0012, ^**^*p=*0.0007, ^***^*p=*7.7× 10^−5, ****^*p=*0.0003 empty vector vs *S. cerevisiae* Hem25p, ^#^*p=*0.0053, ^##^*p=*0.0112, ^###^*p=*0.0004, ^####^*p=*7.0× 10^−5^ empty vector vs *S. pombe* Hem25p, mean ± SD, *n=*3 independent samples).

We next sought to better understand the evolutionary divergence of Hem25p and SLC25A38 function. *HEM25* is highly conserved among opisthokonts. Thus, we used the PANTHER database to compile a list of *HEM25* orthologs from a number of model organisms across this group^44^. We expressed each ortholog in *hem25*Δ yeast and measured their ability to rescue CoQ levels. Full rescue was largely limited to fungal orthologs, with little to no rescue with metazoan orthologs (Fig. 4d). Phylogenic analysis supports this functional split, showing an early evolutionary event resulting in separate fungal and metazoan branches (Fig. 4d)^44^. To validate Hem25p’s role in fungi, we expressed the *Schizosaccharomyces pombe* ortholog in bacteria and measured IPP uptake. Consistent with our rescue experiment, Hem25p from *S. pombe* and *S. cerevisiae—*two highly evolutionarily diverse yeast species—demonstrated similar levels of IPP uptake (Fig. 4e). Collectively, our results show that the mitochondrial glycine carrier Hem25p doubles as an IPP transport in fungi, and will empower subsequent studies to identify IPP transporters in metazoans.

## Discussion

In this work, we identify Hem25p as a mitochondrial IPP transporter. Our results reveal a novel function for this established glycine transporter and propose a model whereby a single carrier, Hem25p, enables the bulk of mitochondrial heme and CoQ biosynthesis. However, even in the absence of Hem25p, heme and CoQ are still produced at low levels (Fig. 2b). This suggests that functional redundancy is inherent among mitochondrial transporters, and may explain why previous large-scale screens for CoQ biosynthetic proteins have missed Hem25p^16,17^. In our targeted screen, we did not find any strong candidates for secondary transporters, despite having screened every known yeast mitochondrial transporter. It is possible that multiple SLC25 transporters each have low level IPP transport capability or that a non-SLC25 protein not included in our screen is the secondary transporter. Compensatory mechanisms may also mask subtle changes when a secondary transporter is disrupted. CoQ biosynthetic proteins have been shown to organize into domains at ER-mitochondria contact sites^45,46^. Given that the mevalonate pathway, which supplies IPP, is partially associated with the ER, one possible function for this co-localization is the transport of substrates and biosynthetic precursors. Thus, it is possible that these domains provide an additional layer of redundancy. Further efforts are needed to dissect the complete mechanisms of isoprenoid pyrophosphate transport.

Hem25p’s ability to support two separate biosynthetic pathways raises the possibility that other mitochondrial transporters carry multiple substrates. Indeed, the yeast Pic2p and its mammalian homolog SLC25A3 were recently shown to transport copper in addition to phosphate—two highly distinct substrates, much like glycine and IPP^47,48^. Thus, in that case, a single carrier enables oxidative phosphorylation by supporting both ATP synthesis and cytochrome *c* oxidase activity. Many metabolites still lack a defined transporter, despite evidence of mitochondrial uptake. The fact that yeast and humans have only 35 and 53 SLC25 carriers, respectively, suggests that functional redundancy is widespread^17^. However, potential contribution from non-SLC25 carriers, many of which remain unidentified, must also be considered. Thus, a complete and accurate definition of mitochondrial transporters and their substrate specificities is still needed. The development of new experimental technologies and screening approaches will accelerate these characterization efforts.

A previous study identified six SLC25 members that genetically interact with *HEM25*, including *FLX1, MTM1*, and *SAM5*^35^. Interestingly, disruption of these three genes resulted in decreased CoQ levels in our screen (Fig. 1b). *FLX1* and *SAM5* encode transporters for FAD and SAM, respectively, both of which are necessary cofactors for CoQ biosynthesis^20–23^. Mtm1p is an uncharacterized mitochondrial transporter that was originally proposed to assist with mitochondrial manganese trafficking. However, it was later suggested that it participates in iron-sulfur cluster biogenesis^24,26^. Cells lacking Mtm1p exhibit elevated mitochondrial manganese levels, which can result in the mismetallation and degradation of Coq7p, the diiron hydroxylase responsible for the penultimate step of CoQ biosynthesis^24,25^. Additionally, *mtm1*Δ cells phenocopy disruptions in the biosynthesis of iron-sulfur clusters—cofactors necessary for electron transfer reactions that enable Coq6p function^7^. Thus, it is possible that the observed decrease of CoQ in *mtm1*Δ is a result of Coq7p mismetallation and degradation, dysfunctional Coq6p, or both. These roles are supported by our CoQ and PPHB measurements and may underlie the observed genetic interactions with *HEM25*. Additional work is needed to confirm the molecular basis of these interactions.

Our results corroborate previous *in vitro* experiments showing that imported IPP is sufficient to generate CoQ’s polyprenyl tail (Fig. 2e)^12,13^. We did not assess whether other isoprenoid pyrophosphates, such as dimethylallyl pyrophosphate (DMAPP), geranyl pyrophosphate (GPP), or farnesyl pyrophosphate (FPP), can be imported. Coq1p, a mitochondrial prenyltransferase, has shown an ability to use GPP and FPP in isolated membrane preparations^49^, but whether these isoprenoids can be transported into mitochondria remains unknown. Moreover, it has been argued that a common cytosolic FPP pool supplies precursors for CoQ and other isoprenoids, however this remains unresolved^50–52^. While our investigation shows that yeast mitochondria can bypass this cytosolic FPP pool by importing and incorporating IPP alone, we cannot exclude the contribution of other isoprenoid pyrophosphate species to CoQ biosynthesis.

Despite being the closest ortholog to Hem25p, SLC25A38 did not transport IPP nor did it functionally rescue CoQ levels in *hem25*Δ (Fig. 4b,c). We also did not identify any reports of CoQ deficiency in patients with SLC25A38 mutations. At first glance, this seems surprising given that both carriers share many of the same key binding site residues (Fig. 3d)^40^. Moreover, we found these conserved residues to be important for Hem25p stability or substrate binding (Fig. 3e,f). However, our experimental and phylogenetic analyses show a clear divergence between fungal and metazoan orthologs, with IPP import limited to fungal orthologs (Fig. 4d). It is likely that the emergence of multicellular and multiorgan animals necessitated a higher level of regulation of heme and CoQ biosynthesis, such that a common carrier was no longer sufficient. Consistent with this model, human *SLC25A38* demonstrates preferential expression in early erythroid cells and both zebrafish orthologs are strongly associated with hematopoiesis and hematopoietic tissues^28,41^. Given that CoQ is produced in nearly every mammalian cell type, we would expect ubiquitous expression from a mammalian isoprenoid transporter. Additional differences in both the heme and CoQ biosynthetic pathways have been identified between yeasts and metazoans^53–55^, and more work is needed to dissect the physiological underpinnings of these differences.

The divergence in Hem25p and SLC25A38 functions implies the existence of a separate metazoan IPP transporter. Such an occurrence would not be unique to IPP, with NAD being the most recent example of a substrate having distantly related fungal and metazoan transporters^56–59^. Our identification of the primary fungal IPP transporter here lays the foundation for future efforts to identify the functional metazoan ortholog.

## Methods

### Yeast strains and cultures

The *S. cerevisiae* haploid strain W303 (MATa leu2 trp1 can1 ura3 ade2 his3) was used. Single (*gene*Δ) and double (*gene*Δ*gene*Δ) deletion strains were generated using standard homologous recombination^60^. Open reading frames were replaced with the kanMX6, His3MX6, or Trp1 cassettes and confirmed by PCR assay^61,62^.

For the targeted screen, cells were grown in YPD media consisting of 1% (w/v) yeast extract (Research Products International, RPI), 2% (w/v) peptone (RPI), and 2% (w/v) dextrose (Fisher). YP media without dextrose were sterilized by automatic autoclave. Glucose was sterile filtered (0.22 *μ*m pore size, VWR) and added to sterile YP prior to use. For all other cultures, synthetic complete (SC) or synthetic dropout (SD) media was used, containing yeast nitrogen base (YNB) with ammonium sulfate and without amino acids (US Biological), the corresponding drop-out mix (US Biological), and the indicated carbon source. pABA^−^ media contained YNB without amino acids and without pABA (Formedium), Complete Supplement Mixture (Formedium), and the indicated carbon source. All synthetic yeast media were sterile filtered (0.22 *μ*m pore size, VWR) prior to use. Where indicated, ALA or glycine were added to the media prior to sterile filtration.

For measurements in respiration^30^, starter cultures (3 mL, SC or SD, 2%D) were inoculated with individual colonies and incubated (30 ºC, 230 rpm, 14–16 h). Cell density was measured at OD600 and converted to cells/mL (1 OD = 1 × 10^7^ cells/mL). Respiratory media (100 mL, SC or SD, 0.1%D, 3%G) in a sterile 250 mL Erlenmeyer flask were inoculated with 2.5 × 10^6^ yeast cells. Samples were incubated (30 ºC, 230 rpm) for 25 h, a time point that corresponds to early respiratory growth.

### Lipid extraction

Lipid extractions were performed essentially as described previously^63^. 1×10^8^ cells were harvested by centrifugation (4,000 ×*g*, 5 min, RT). The supernatant was removed and the cells washed with water (600 *μ*L). Cells were pelleted again (15,000 × *g*, 30 s, RT) and the supernatant removed. Cell pellets were either used immediately for lipid extraction or snap-frozen in LN2 and stored at −80 °C until analysis. Frozen pellets were thawed on ice prior to extraction. To extract lipids from whole cells, 150 mM KCl (50 *μ*L) was added to each sample, followed by ice-cold methanol (600 *μ*L; with 1 *μ*M CoQ8 as an internal standard). Glass beads (100 *μ*L; 0.5 mm; BioSpec) were then added and the samples were vortexed (10 min, 4 ºC) to lyse the cells. Ice-cold petroleum ether (400 °L; Sigma) was added to extract lipids, and the samples were vortexed again (3 min, 4 °C). Samples were centrifuged (1000 × *g*, 3 min, RT) and the top petroleum ether layer was collected in a new tube. The petroleum ether extraction was repeated a second time, with the petroleum ether layer from the second extraction combined with that from the first. The extracted lipids were dried under argon before being resuspended in 2-propanol (50 *μ*L) and transferred to an amber glass vial (Sigma; QSertVial, 12 × 32 mm, 0.3 mL).

### Targeted yeast genetic screen

Yeast from a −80 °C glycerol stock were struck on to YPD plates and incubated (30 °C, 230 rpm, 48 h). Starter cultures of YPD (3 mL) were inoculated with individual colonies and grown overnight (30 °C, 14–16 h). Overnight cultures were diluted to an OD600 = 0.2 (3 mL) and incubated until OD600∼1 (30 °C, 230 rpm, 4-5 h), corresponding to mid-log phase. Cultures were harvested by centrifugation (4,000 x *g*, 5 min, RT), washed once with water, and transferred to a 1.5 mL microcentrifuge tube. Cells were centrifuged again (15,000 x *g*, 30 s, RT), snap frozen in LN2, and stored at −80 °C until analysis. Lipids were extracted from cell pellets as described above and CoQ measurements were then performed by HPLC-ECD.

### Targeted HPLC-ECD for CoQ

Extracted lipids were resuspended in 2-propanol (50 *μ*L) and transferred to an amber vial. Sodium borohydride (2 *μ*L of 10 mM in 2-propanol) was added to each vial, followed by brief vortexing and incubation (10 min, RT) to reduce CoQ. Methanol (50 *μ*L) was then added to each sample to remove excess sodium borohydride and the vials were flushed with argon gas. CoQ measurements were performed using reverse-phase high-pressure liquid chromatography with electrochemical detection (HPLC-ECD)^63^. Separation was performed using C18 column (Thermo Scientific, Betasil C18, 100 × 2.1 mm, particle size 3 *μ*m) at a flow rate of 0.3 mL/min with a mobile phase of 78% methanol, 10% 2-propanol, 10% acetonitrile, and 2% ammonium acetate (1 M, pH 4.4). Electrochemical detection was performed using an ECD detector (Thermo Scientific ECD3000-RS) containing a 6020RS omni Coulometric Guarding Cell (E1) set to −200 mV and two 6011RS ultra Analytical Cells (E2 and E3) set to 600 mV. CoQ measurements were made on the analytical E2 electrode. For each experiment, CoQ_6_ and CoQ_8_ standards (Avanti) were prepared in the same manner as the experimental samples and injected to generate a standard curve. Peak areas were quantified using Chromeleon 7.2.10 software (Thermo), normalized to the CoQ_8_ internal standard, and converted to absolute values using the standard curve. CoQ_6_ levels were further normalized per mg of wet pellet weight.

### Heme measurements

Yeast total heme measurements were performed using previously described methods with slight modifications^64,65^. Cells were grown under respiratory conditions as described previously and supplemented with glycine or ALA as indicated. 1 × 10^8^ cells were harvested by centrifugation (4,000 × *g*, 5 min, RT), washed with water, and centrifuged again (15,000 × *g*, 30 s, RT). Pellets were snap frozen in LN2 and stored at −80 °C. Frozen pellets were thawed on ice before being resuspended in 500 *μ*L oxalic acid (20 mM) and incubated in a closed box (16-24 h, 4 °C). 500 *μ*L oxalic acid (2 M, preheated to 50 °C) were added to each sample, and each sample was divided in half into two 1.5 mL amber microcentrifuge tubes. One half of each sample was heated (95-100 °C, 30 min) while the other was incubated at RT. Each tube was then centrifuged (16,000 × *g*, 2 min) and 200 *μ*L of the supernatant were loaded into a 96-well black-bottom plate (Greiner). Heme fluorescence was measured on a Cytation 3 plate reader (BioTek) with excitation at 400 nm and emission at 620 nm. The fluorescence from the unheated sample was subtracted from the corresponding heated sample.

### Quantitative PCR for *COQ* gene expression

Respiring yeast were cultured and harvested as described in “*Yeast strains and cultures*” for heme measurements. Total RNA was extracted using the MasterPure Yeast RNA Purification Kit (Lucigen). First strand cDNA synthesis was performed using the EasyScript Plus cDNA Synthesis Kit (Lamda Biotech) using Oligo(dT) primers and 1 *μ*g of RNA. qPCR was performed using the following reaction: 10 *μ*L *Power* SYBR Green PCR Master Mix (Thermo), 2.5 *μ*L cDNA (diluted 1:20), and 250 nM forward and reverse primers. Primers amplifying the yeast *COQ2, COQ3, COQ4, COQ5, COQ6, COQ7, COQ8, COQ9*, and *ACT1* (reference) genes were used. For the qPCR cycle, an initial 2 min incubation at 50 °C was followed by 10 min denaturing at 95 °C. Then 40 cycles of 95 °C for 15 s and 60 °C for 1 min were performed. RNA abundance was calculated using the ΔΔC*t* method.

### LC-MS/MS proteomics

#### Yeast growth

Yeast cultures were grown as described previously for respiration. 1 × 10^8^ cells were harvested, snap frozen in LN2, and stored at −80 °C.

#### Lysis and Digestion

Yeast pellets were removed from −80 °C conditions and resuspended in lysis buffer (6 M guanidine hydrochloride, 100 mM Tris). The samples were then boiled at 100°C for 5 minutes and sonicated in a bath sonicator (Qsonica) with a 5-minute-long program of 20 seconds on, 10 seconds off. Methanol was added to each sample to 90% concentration, and the samples were centrifuged at 10,000 × g for 5 minutes to precipitate proteins. After precipitation, the supernatant was discarded from each sample, and the protein pellets were allowed to air dry for 7 minutes. The dried pellets were resuspended in digestion solution (8 M urea, 10 mM TCEP, 40 mM CAA, 100 mM Tris), and the samples were sonicated in the bath sonicator with the same program as above to facilitate resolubilization. LysC (Wako Chemicals) was added to each resolubilized sample in an estimated 50:1 protein/enzyme ratio. The samples were incubated on a rocker at room temperature for 4 hours before being diluted fourfold with 100 mM Tris. Trypsin (Promega) was added to each sample in an estimated 50:1 protein/enzyme ratio before they were incubated on a rocker at room temperature for 14 hours. Each sample was finally acidified with TFA to pH of 2, desalted by solid phase extraction cartridges (Phenomenex), and dried under vacuum (Thermo Scientific).

#### LC-MS/MS Proteomics Data Acquisition

Peptides were resuspended in 0.2% formic acid, and the concentration of each sample was determined from a NanoDrop One spectrophotometer (Thermo Scientific). The samples were then prepared in autosampler vials and loaded onto a 75 *μ*m i.d. x 360 *μ*m o.d. capillary column (New Objective) that was packed in-house^66^ with 1.7 *μ*m BEH C18 particles and held at 50 °C throughout the analysis. Separations were performed with a Dionex UltiMate 3000 nano HPLC system (Thermo Scientific). The mobile phases were 0.2% formic acid in water (A) and 0.2% formic acid in 80% ACN (B). The peptides were analyzed by an Orbitrap Eclipse (Thermo Scientific) with the following parameters: Orbitrap MS1 resolution of 240,000; MS1 automatic gain control target of 1 × 10^6^; MS1 maximum injection time of 50 ms; MS1 scan range of 300-1500 *m/z*; dynamic exclusion of 20 ms; advanced peak determination^67^ toggled on; MS2 isolation window of 0.7 *m/z*; MS2 collision energy of 25%; ion trap MS2 resolution setting of turbo; MS2 automatic gain control target of 3 × 10^4^; MS2 maximum injection time of 14 ms; and MS2 scan range of 150-1200 *m/z*.

#### Data Analysis

MaxQuant^68^ (version 1.5.2.8) was used to process proteomics raw files against a database of reviewed yeast proteins plus isoforms from UniProt (downloaded 11/20/2019). Label-free quantification^69^ was used for relative quantification. Match-between-runs was enabled. Data were visualized and analyzed using the Argonaut^70^ platform.

#### Data Availability

Proteomics raw files have been deposited to the MassIVE repository, with accession number MSV000089127. Reviewers may access the raw files through the following link with the password “proteomics”: ftp://MSV000089127@massive.ucsd.edu.

### Targeted LC-MS/MS for PPHB and CoQ

#### Yeast growth

For measurements in respiration, yeast cells were grown and harvested as described previously. Yeast pellets were snap frozen in LN_2_ and stored at −80 °C.

For *de novo* PPHB and CoQ experiments, starter cultures (3 mL, SC, 2%D, 300 *μ*M ALA) were inoculated with individual colonies and incubated (30 °C, 230 rpm, 18 h). OD_600_ was measured, and 2.5 × 10^7^ cells were diluted into 50 mL of pABA^−^ media (2%D, 300 *μ*M ALA) in a sterile 250 mL Erlenmeyer flask. Cells were incubated (30 °C, 230 rpm) until OD_600_ ∼ 1.5-2 to deplete pABA in the cells. 7.5 × 10^8^ cells were centrifuged (3,000 × *g*, 5 min, RT) and resuspended in 50 mL labeled pABA^−^ media (3%G, 300 *μ*M ALA, 50 *μ*M ^13^C6-4-HB). Immediately after the media swap, and at 2, 4, and 6 h after, 1 ×10^8^ cells were harvested by centrifugation (3,000 × *g*, 5 min, 4°C) and washed with water. Cells were pelleted again (15,000 ×*g*, 30 s, 4 °C), snap frozen in LN2, and stored at −80 °C.

#### LC-MS/MS Lipidomics Data Acquisition

Frozen pellets were thawed on ice and lipid extraction was performed as described previously. A Vanquish Horizon UHPLC system (Thermo Scientific) connected to an Exploris 240 Orbitrap mass spectrometer (Thermo Scientific) was used for targeted LC-MS analysis. A Waters Acquity CSH C18 column (100 mm × 2.1 mm, 1.7 *μ*m) was held at 35°C with the flow rate of 0.3 mL/min for lipid separation. A Vanquish binary pump system was employed to deliver mobile phase A consisted of 5 mM ammonium acetate in ACN/H_2_O (70/30, v/v) containing 125 *μ*L/L acetic acid, and mobile phase B consisted of 5 mM ammonium acetate in IPA/ACN (90/10, v/v) containing 125 *μ*L/L acetic acid. The gradient was set as follows: B was at 2% for 2 min and increased to 30% over the next 3 min, then further ramped up to 50% within 1 min and to 85% over the next 14 min, and then raised to 99% over 1 min and held for 4 min, before re-equilibrated for 5 min at 2% B. Samples were ionized by a heated ESI source with a vaporizer temperature of 350°C. Sheath gas was set to 50 units, auxiliary gas was set to 8 units, sweep gas was set to 1 unit. The ion transfer tube temperature was kept at 325°C with 70% RF lens. Spray voltage was set to 3,500 V for positive mode and 2,500 V for negative mode. The targeted acquisition was performed with both tMS_2_ (targeted MS2) mode and tSIM (targeted selected ion monitoring) mode in the same injection: tMS_2_ mode was for measuring CoQ_6_ (*m/z* 591.4408), [13]C6-CoQ6 (*m/z* 597.4609) and CoQ_8_ (*m/z* 727.5660, internal standard) in positive polarity at the resolution of 15,000, isolation window of 2 *m/z*, normalized HCD collision energy of either 40% or stepped HCD energies of 30% and 50%, standard AGC target and auto maximum ion injection time; tSIM mode was for measuring PPHB_6_ (*m/z* 545.4000), [13]C6-PPHB_6_ (*m/z* 551.4201), and [13]C12-PPHB6 (*m/z* 557.4403) in negative polarity at the resolution of 60,000, isolation window of 2 *m/z*, standard AGC target and auto maximum ion injection time.

#### Data Analysis

Targeted quantitative analysis of all acquired compounds was processed using TraceFinder 5.1 (Thermo Scientific) with the mass accuracy of 5 ppm. The result of peak integration was manually examined.

### Mitochondrial isolation

Starter cultures (3 mL, SC or SD, 2%D) were inoculated with individual colonies and incubated (30 °C, 230 rpm, 14-16 h). 2 × 10^8^ cells were diluted into 1 L SC 2%D in a 5 L Erlenmeyer flask and incubated (30 °C, 230 rpm) for 14-18 to a final OD ∼6. Mitochondria were isolated as previously described^71^. Cells were harvested by centrifugation (3000 *g*, 5 min, RT), washed with water, and centrifuged again (3,000 × *g*, 5 min, RT). The wet pellet weight of the cells was determined. Cells were resuspended in 2 ml/g DTT buffer (100 mM Tris-H_2_SO_4_, 10 mM DTT, pH 9.4) and shaken slowly (30 ºC, 80 rpm, 20 min). Cells were pelleted, washed once with 7 ml/g Zymolyase buffer (1.2 M sorbitol, 20 mM potassium phosphate, pH 7.4), and resuspended in 7 mL/g Zymolyase buffer with 3 mg/g Zymolyase 20T (Fisher) to generate spheroplasts. Yeast were shaken slowly (30 ºC, 80 rpm) for 30 min before being pelleted and washed with 7 mL/g Zymolyase buffer. Pellets were resuspended in 6.5 mL/g ice-cold homogenization buffer (0.6 M sorbitol, 10 mM Tris-HCl, 1 mM PMSF, 0.2% (w/v) fatty acid-free BSA, pH 7.4). Spheroplasts were homogenizated using 15 strokes of a tight-fitting glass-Teflon homogenizer and diluted twofold with homogenization buffer. The homogenate was centrifuged (1500 × *g*, 5 min, 4 °C) to pellet cell debris and nuclei. The supernatant was centrifuged (4,000 × *g*, 5 min, 4 °C) to pellet additional debris, and the supernatant centrifuged again (4,000 × *g*, 5 min, 4 °C). Mitochondria were isolated by centrifuging the supernatant (12,000 × g, 15 min, 4 °C) and resuspended in SEM (250 mM sucrose, 1 mM EDTA, 10 mM MOPS-KOH, pH 7.2) or SEP (0.6 M sorbitol, 1 mM EGTA, 50 mM potassium phosphate, pH 7.4), where indicated. Mitochondrial protein content was quantified by BCA protein assay (Thermo).

### Mitochondrial PPHB synthesis

Isolated mitochondria were resuspended in SEP at a concentration of 8 mg/mL, kept on ice, and used within 4 h of isolation without freezing. PPHB synthesis was carried out using a modified protocol^12^. To begin the assay, 250 *μ*L mitochondria (2 mg per assay) were added to 750 *μ*L of 1.3X substrate buffer (0.6 M sorbitol, 50 mM potassium phosphate, 1 mM EGTA, 9.1 mM MgCl_2_, 1.3 *μ*M 4-HB, 13.3 *μ*M [1,2-^13^C]-IPP (Cambridge Isotope Laboratories), pH 7.4). The final concentrations in the reaction were the following: 0.6 M sorbitol, 50 mM potassium phosphate, 1 mM EGTA, 7 mM MgCl_2_, 1 *μ*M 4-HB, 10 *μ*M [1,2-^13^C]-IPP, and 2 mg mitochondria, pH 7.4. Reactions were carried out at 30 ºC in a 5 mL tube (Eppendorf) with constant shaking (500 rpm). At the indicated time points, 300 *μ*L of the reaction were removed and immediately extracted for lipids. 300 *μ*L ice-cold methanol containing 1 *μ*M CoQ_8_ as an internal standard were added to each sample, followed by 400 *μ*L ice-cold petroleum ether. Samples were vortexed (10 min, 4 °C) and centrifuged (3,000 x *g*, 3 min, 4 °C). The top petroleum ether layer was collected, and the extraction repeated again with another 400 *μ*L petroleum ether. The petroleum ether layers were pooled for each sample, dried under argon, and stored at -80 ºC. Lipids were resuspended in 2-propanol (50 *μ*L) and transferred to amber glass vials for targeted LC-MS/MS. Following the third time point, 50 *μ*L of the remaining reaction was diluted with 2X SDS sample buffer for immunoblot analysis.

### Bacterial uptake assays

Bacterial expression of mitochondrial carriers and whole-cell uptake assays were performed as described previously with slight modifications^36,37^. MBP-tagged fusion proteins were generated by combining the bacterial maltose binding protein (MBP) containing the MalE signal peptide, a short linker containing a thrombin cleavage site, and the mitochondrial carrier gene. Fusion constructs were synthesized as gBlock gene fragments (IDT) and cloned into the pET-21b expression vector using the restriction enzymes NdeI and XhoI. Inserts were confirmed by sequencing. Mutants were generated by site-directed mutagenesis (NEB).

Expression of MBP-tagged carriers was carried out in E. *coli* C43(DE3) cells (Biosearch Technologies)^72^. Single colonies of transformed cells were used to inoculate 4 mL of LB media containing 100 mg/L ampicillin. Cultures were incubated overnight (37 °C, 230 rpm) before being diluted 1:100 in 50 mL LB media containing 100 mg/L ampicillin. Refreshed cultures were incubated (37 °C, 230 rpm) until the OD600=0.5-0.6, at which point protein expression was induced with 0.1 mM IPTG. Cells were induced overnight (20 ºC, 230 rpm, 14-16 h).

Induced cells were collected by centrifugation (4,000 × *g*, 10 min, 4 °C), washed once with ice-cold KPi (50 mM potassium phosphate, pH 7.4), and centrifuged again (4,000 × *g*, 10 min, 4 °C). Cells were resuspended in ice-cold KPi to a cell density of OD_600_=10 and placed on ice until the start of the assay.

To start the assay, 200 *μ*L of cells were added to 200 *μ*L of 2X substrate buffer (2 mM MgCl_2_ and radiolabeled substrate at double the final concentration in KPi). Assays were carried out at room temperature. [1-^14^C]-IPP and [*phenyl*-^14^C]-4-HB were purchased from American Radiolabeled Chemicals. At each time point, 100 *μ*L of the assay mixture were removed and filtered under vacuum to separate the cells from the incubation media. To reduce non-specific binding of the substrate to the filter, filtration was carried out using 0.22 *μ*m mixed cellulose ester (MCE) filters (Millipore) for IPP uptake assays and 0.2 *μ*m Whatman Nuclepore track-etched hydrophilic membrane filters (Cytiva) for 4-HB uptake assays^39^. Following filtration, filters were washed with 5 mL of ice-cold KPi before being placed in a 7 mL scintillation vial (Fisher). 5 mL of Ultima Gold MV liquid scintillation cocktail (PerkinElmer) were added to each vial before being analyzed by liquid scintillation counting. Counts per minute were converted to disintegrations per minute (DPM) using the counting efficiency of the counter.

### CoQ rescue experiments

The yeast expression vector p416 GPD was modified to contain the endogenous *HEM25* promoter. PCR was used to amplify a 744 bp segment directly upstream of the *HEM25* gene from yeast genomic DNA. This segment, containing the endogenous promoter, was then cloned into the SacI-BamHI sites of the p416 GPD vector, yielding a p416 vector containing the *<u>H</u>EM25* <u>e</u>ndogenous <u>p</u>romoter (p416 HEP). The *HEM25* gene was amplified from yeast genomic DNA and cloned into the BamHI-XhoI sites of the p416 GPD or the p416 HEP vectors. The PCR primers were designed so that a short linker and a FLAG tag was added to the end of the open reading frame. *HEM25-*FLAG mutants were synthesized as gBlocks (IDT) and cloned into the BamHI-XhoI sites of the p416 GPD or the p416 HEP vectors. We were not able to PCR amplify p416, thus SDM could not be performed. The sequences of the cloned *HEM25* promoter and the inserts were confirmed by sequencing. Verified constructs were transformed into *hem25*Δ yeast using the LiAc/SS carrier DNA/PEG method^73^. For controls, empty vectors were also transformed into WT W303, *coq6*Δ, *hem25*Δ yeast. Respiratory growth and targeted lipidomics were performed as described above.

*HEM25* orthologs were synthesized as codon-optimized gBlocks (IDT) and cloned into the BamHI-XhoI sites of the p416 ADH vector. Constructs were sequenced and transformed into *hem25*Δ yeast. Fermentative growth and targeted lipidomics were performed as described above.

### Sequence alignments and bioinformatics analysis

Hem25p orthologs were curated from the PANTHER database^44^. Sequences were aligned using Clustal Omega in Jalview^74,75^. The phylogenetic tree was generated using Jalview and FigTree. The structure of Hem25p was predicted using AlphaFold and visualized using PyMol 2.0^76,77^.

### Immunoblotting

#### Antibodies

Primary antibodies for this study include anti-FLAG (Sigma F1804, 1:2500-1:5000), anti-Act1 (Abcam ab8224, 1:5000), anti-Coq1 (custom made at Genscript, 1:2000), anti-Vdac1 (Abcam ab110326, 1:4000), and Anti-MBP (NEB E8032S, 1:10000). Secondary antibodies include IRDye 680RD goat anti-rabbit (LI-COR 926-68071, 1:15000), IRDye 680RD goat anti-mouse (LI-COR 926-68070, 1:15000), IRDye 800CW (LI-COR 926-32210, 1:15000), and anti-mouse IgG HRP-linked (Cell Signaling Technology #7076, 1:2000-1:100000).

#### Whole yeast cell lysates

Protein lysates from whole yeast cells were prepared as described^78^. Yeast pellets were lysed using 150 *μ*L of lysis buffer (2 M NaOH, 1 M β-mercaptoethanol) for 10 min on ice with periodic vortexing. 150 *μ*L of 50% TCA was added, and the samples were incubated on ice for 10 min with periodic vortexing to precipitate proteins. The samples were centrifuged (14,000 × *g*, 2 min, RT) and the supernatant discarded. The remaining pellet was washed with 1 mL acetone before being pelleted again (14,000 × *g*, 2 min, RT). The pellet was left to air dry before being resuspended in 120 *μ*L of 0.1 M NaOH. Protein concentrations were determined by BCA protein assay (Thermo) and diluted twofold with 2X SDS sample buffer.

30 *μ*g of protein were loaded onto NuPAGE 4-12% Bis-Tris Gels (Thermo) and separated (200 V, 35 min). Proteins were transferred to a PVDF membrane (Sigma) and blocked with 5% non-fat dry milk in TBST (1 h, RT). Membranes were then probed with primary anti-FLAG (Sigma F1804, 1:2500) antibodies diluted in 5% (NFDM) in TBST (overnight, 4 °C). Membranes were washed three times with TBST and then probed with secondary antibodies diluted in 5% NFDM in TBST (1 h, RT). Membranes were washed three timed with TBST and developed by enhanced chemiluminescence (ECL). For the blotting of cells harboring constructs containing the endogenous *HEM25* promoter, the SuperSignal West Atto substrate (Thermo, A38554) was used. Blotting for cells harboring constructs containing the constitutive GPD promoter used the SuperSignal West Dura substrate (Thermo, 34075). Developed membranes were imaged on a ChemiDoc system (Bio-Rad) before being stripped (Thermo, 46430). Stripped membranes were blocked in 5% NFDM in TBST (1 h, RT) before being probed with anti-Act1 (Abcam ab8224, 1:5000) loading control antibody (1 h, RT). Membranes were washed three times with TBST before being probed with the secondary antibody diluted in 5% NFDM (1 h, RT). Membranes were washed three times before being developed using the SuperSignal West Dura substrate. Membranes then were imaged using the ChemiDoc.

#### Isolated mitochondria

Samples were collected and prepared as described in “Mitochondrial PPHB synthesis.” 15 *μ*g of protein were loaded onto a NuPAGE 4-12% Bis-Tris Gel (Thermo) and separated (200 V, 35 min). Proteins were transferred to a PVDF membrane (Sigma), cut between the expected sizes of Coq1p and VDAC, and blocked with 4% non-fat dry milk in TBST (1 h, RT). The membranes were then probed with primary antibodies diluted in 4% NFDM (overnight, 4 °C). Membranes were washed three times with TBST before being probed with secondary antibodies (1 h, RT). Membranes were washed three times with TBST before being imaged on a LI-COR Odyssey CLx Imaging System and analyzed with LI-COR Image Studio Software (version 5.2.5).

#### E. coli membranes

Approximately 40 mL*OD of induced cells were collected (4000 × *g*, 15 min, 4 °C), the supernatant removed, snap frozen in LN_2_, and stored at − 80 °C until blotting. Frozen pellets were thawed on ice and resuspended in 2 mL ice-cold lysis buffer (10 mM Tris pH 7 containing protease inhibitors). Cells were sonicated until clear (Branson Ultrasonics, 45% amplitude, 5 s pulse, 50% duty cycle, microtip). Lysed cells were diluted with ice-cold lysis buffer to a final volume of 9 mL. Unlysed cells and insoluble material were pelleted by centrifugation (27,000 × *g*, 15 min, 4 °C). The supernatant was transferred to a 14 mL tube (Beckman 344060) and ultracentrifuged to pellet membranes (150,000 × *g*, 1 h, 4 °C). The supernatant was removed, and the membrane pellet was resuspended in 75 *μ*L ice-cold lysis buffer. Protein concentrations were determined by BCA protein assay (Thermo) and the sample diluted twofold in 2X SDS sample buffer.

5 *μ*g of protein were loaded onto a NuPAGE 4-12% Bis-Tris Gel (Thermo) and separated (150 V, 1 h). Proteins were transferred to a PVDF membrane (Sigma) and blocked with 5 % NFDM in TBST. The membrane was then probed with the primary anti-MBP antibody (NEB E8032S, 1:10000) diluted in 5% NFDM (overnight, 4 °C). The membrane was washed three times with TBST and probed with secondary antibodies (1 h, RT) diluted in 5% NFDM in TBST. The membrane was washed three times with TBST before being imaged on a LI-COR Odyssey CLx Imaging System.

### Statistical analysis

All experiments were performed in at least biological triplicate, unless otherwise stated. All results are presented as the arithmetic mean ± standard deviation (SD). Statistical analysis were performed in Prism 9.0 (GraphPad Software) or in Microsoft Excel. *P* values were calculated using unpaired, two-sided Student’s *t*-test with values less than 0.05 considered significant. For fold changes, the average of the control or wild-type replicates was set to 1, with the other samples normalized accordingly.

## Acknowledgements

We would like to thank members of the Pagliarini lab and K. Henzler-Wildman for helpful discussions and insights throughout this project. This work was supported by NIH R35 GM131795 (D.J.P.), P41 GM108538 (J.J.C. and D.J.P.), F31 AG064891 (J.T.), T32 GM140935 (J.T.), T32 DK007120 (S.W.R.), and the BJC Investigator Program (D.J.P.)

## Data and materials availability

All data are available in the main text or the supplementary materials. Other relevant data and materials are available from the corresponding author upon reasonable request.

## Author contributions

Conceptualization: JT, DJP

Methodology: JT, RMG, SWR, ZF, KAO, DJP

Formal analysis: JT, RMG, SWR, ZF, LKM, ES, KAO

Investigation: JT, RMG, SWR, ZF, LKM, ES, KAO

Funding acquisition: DJP

Supervision: JJC, DJP

Writing – original draft: JT, DJP

Writing – review & editing: JT, RMG, SWR, ZF, LKM, ES, KAO, JJC, DJP

## Supplementary information

Supplementary information is available for this paper.

**Correspondence and requests for materials** should be addressed to D.J.P.

## Supplementary figures

**Supplementary Fig. 1.**
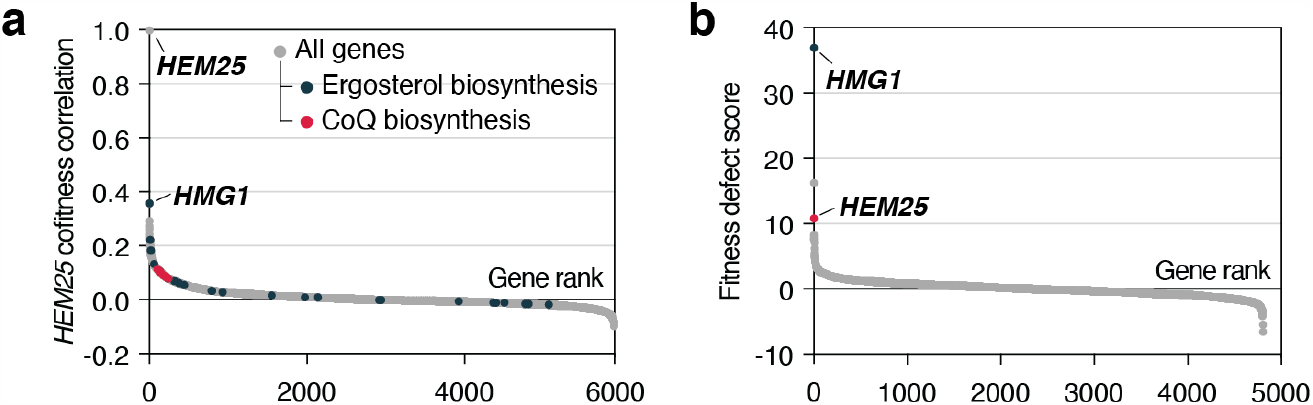
Chemical genomic screens link *HEM25* to the mevalonate pathway. **a**, Gene fitness correlations with *HEM25*. Genes involved with ergosterol and CoQ biosynthesis are highlighted. **b**, Fitness ranking of *gene*Δ strains in the presence of 78.68 *μ*M atorvastatin. Raw data for (**a**) and (**b**) are from the HIPHOP chemogenomics database^31^. Fitness defect scores reflect the homozygous deletion profiles for atorvastatin (compound SGTC_2648).

**Supplementary Fig. 2.**
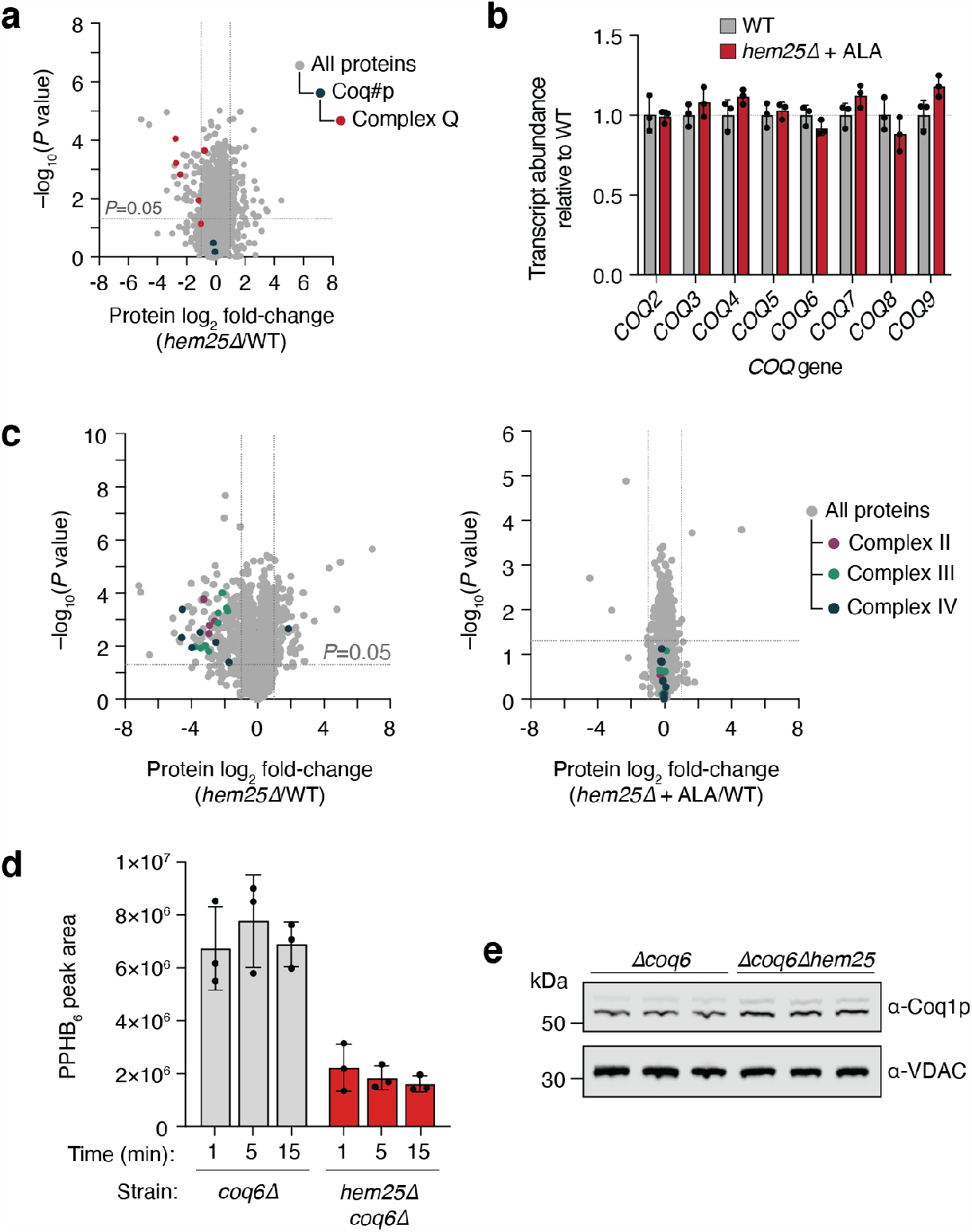
Hem25p drives CoQ biosynthesis independently of heme. **a**, Relative protein abundances of *hem25*Δ cells compared to WT cells verses statistical significance. CoQ biosynthetic proteins and Complex Q proteins are highlighted. Raw data from the Y3K dataset^30^ (respiration-RDR condition) are displayed as the mean from 3 independent samples with two-sided Student’s *t-*test used. **b**, Relative protein abundances of *hem25*Δ cells ± ALA supplementation compared to WT cells without ALA supplementation verses statistical significance. Components of the electron transport chain are highlighted (mean, *n*=3 independent samples, two-sides Student’s *t-*test). **c**, Relative transcript abundances of CoQ biosynthetic genes from *hem25*Δ with ALA supplementation compared to WT cells without ALA supplementation (mean, *n*=3 independent samples). **d**, Normalized abundance of unlabeled PPHB levels in isolated *coq6*Δ and *coq6*Δ*hem25*Δ mitochondria (mean ± SD, *n*=3 independent samples). **e**, Coq1p levels in isolated mitochondria from each replicate, assessed by immunoblotting.

**Supplementary Fig. 3.**
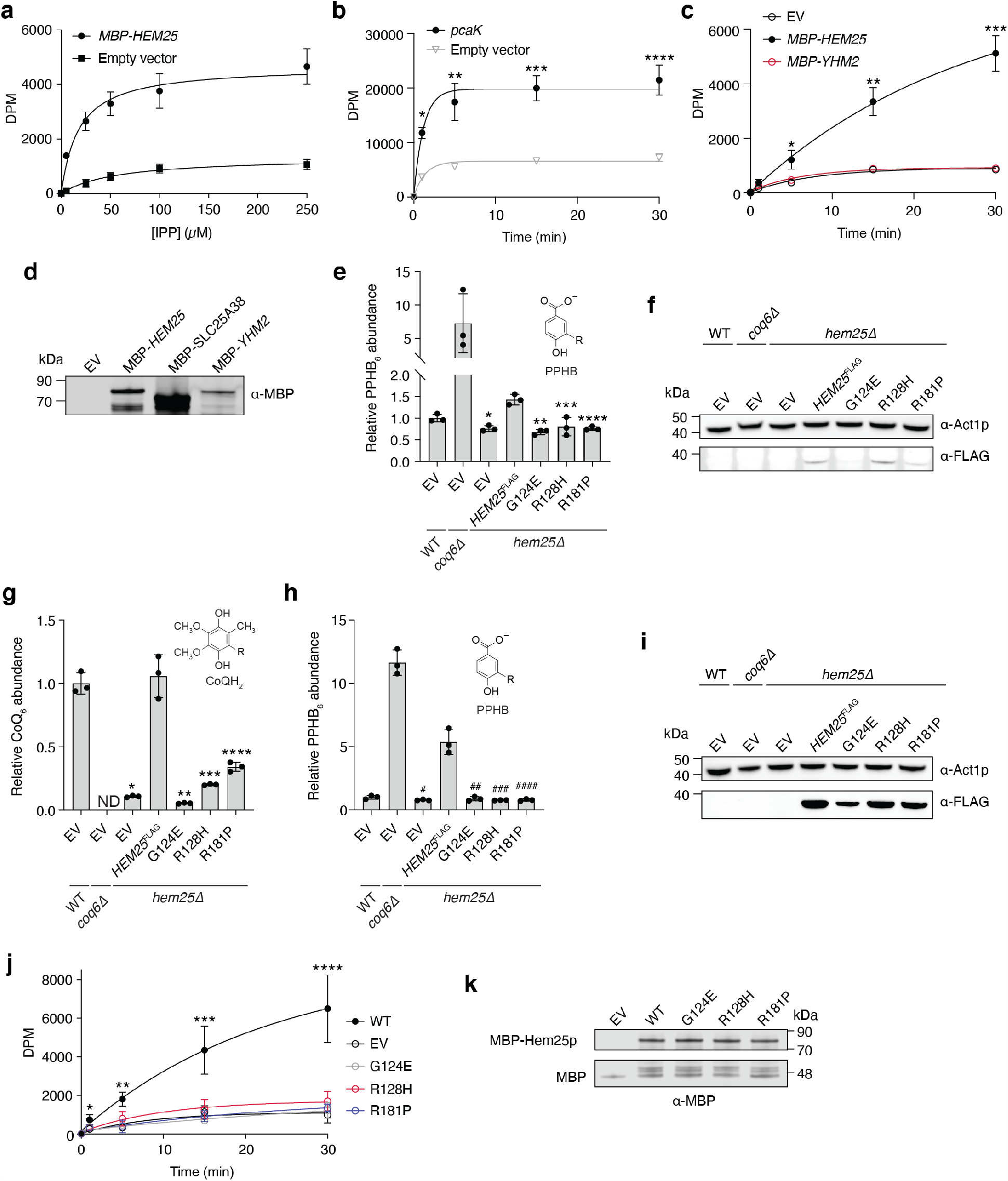
Hem25p enables IPP import into bacteria. **a**, Steady-state kinetics of [1-^14^C]-IPP uptake in cells expressing MBP-Hem25p or the empty vector. Raw DPM values represent the mean ± SD of 3 independent samples. **b**, Time course of 50 *μ*M [*phenyl*-^14^C]-4-HB uptake by *E. coli* cells expressing the *Pseudomonas putida* PcaK or the empty vector (^*^*p=*0.0002, ^**^*p=*0.0038, ^***^*p=*0.0006, ^****^*p=*0.0009 empty vector vs *pcaK*, mean ± SD from 3 independent samples) **c**, Time course of 50 *μ*M [1-^14^C]-IPP uptake by *E. coli* cells expressing MBP-Hem25p, MBP-Yhm2p, or the empty vector (^*^*p=*0.0147, ^**^*p=*0.0011, ^***^*p=*0.0003 empty vector vs *MBP-HEM25*, mean ± SD, *n=*3 independent samples). **d**, Immunoblot for MBP-Hem25p, MBP-SLC25A38, and MBP-Yhm2p in *E. coli* membranes preparations. **e**, Relative PPHB abundances in *hem25*Δ yeast carrying WT and mutant *HEM25*^FLAG^ constructs under the control of the endogenous *HEM25* promoter. Levels are relative to WT yeast carrying the empty expression vector (^*^*p=*0.0011, ^**^*p=*0.0006, ^***^*p=*0.0116, ^****^*p=*0.0009 *HEM25*^FLAG^ vs mutants or empty vector, mean ± SD from 3 independent samples). **f**, Immunoblot of *hem25*Δ cells expressing WT and mutant Hem25p-FLAG under the control of the endogenous *HEM25* promoter. **g, h**, Relative (**f**) CoQ and (**g**) PPHB abundances in *hem25*Δ yeast carrying WT and mutant *HEM25*^FLAG^ constructs under the control of the constitutive GPD promoter. Levels are relative to WT yeast carrying the empty expression vector (^*^*p=*0.0006, ^**^*p=*0.0005, ^***^*p=*0.0009, ^****^*p=*0.0019, ^#^*p=*0.0012, ^##^*p=*0.0014, ^###^*p=*0.0012, ^####^*p=*0.0012 *HEM25*^FLAG^ vs mutants or empty vector, mean ± SD from 3 independent samples); **i**, Immunoblot of *hem25*Δ cells expressing WT and mutant Hem25p-FLAG under the control of the constitutive *GPD* promoter. **j**, Time course of 50 *μ*M [1-^14^C]-IPP uptake by *E. coli* cells expressing WT MBP-Hem25p, mutant MBP-Hem25p, or the empty vector (^*^*p=*0.0429, ^**^*p=*0.0021, ^***^*p=*0.0124, ^****^*p=*0.006 empty vector vs WT, mean ± SD, *n=*3 independent samples). **k**, Immunoblot for WT and mutant MBP-Hem25p in *E. coli* membranes preparations.

**Supplementary Fig. 4.**
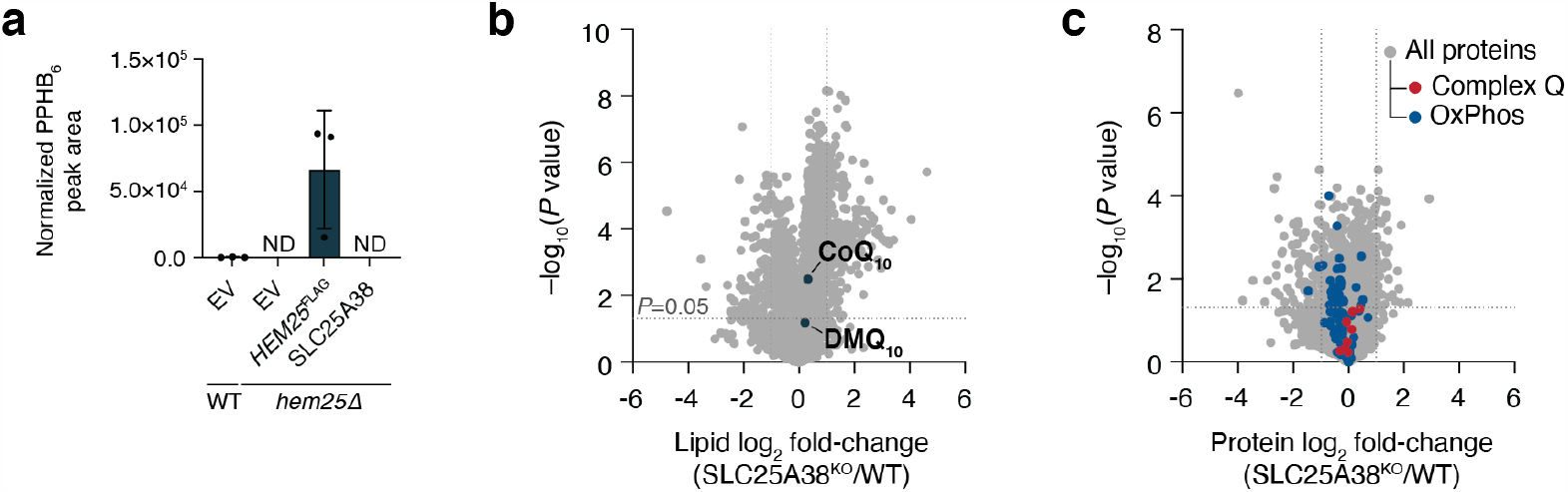
SLC25A38 does not contribute to CoQ biosynthesis. **a**, Normalized PPHB abundance in *hem25*Δ cells expressing Hem25p-FLAG or SLC25A38. (mean ± SD from 3 independent samples) **b**, Relative lipid abundances in *hem25*Δ yeast compared to WT verses statistical significance with CoQ_10_ and the CoQ_10_ biosynthetic intermediate demethoxy-coenzyme Q (DMQ_10_,) highlighted. **c**, Relative protein abundance SLC25A38^KO^ cells compared to WT cells verses statistical significance with CoQ-related (COQ3-COQ9) and OXPHOS-related proteins highlighted. For panels (**b**) and (**c**), raw lipidomic and proteomic data, respectively, from the MITOMICS resource^43^ are displayed as the mean from 3 independent samples with two-sided Welch’s *t-*test used.

## Supplelemtary Tables

**Supplementary Table 1:**
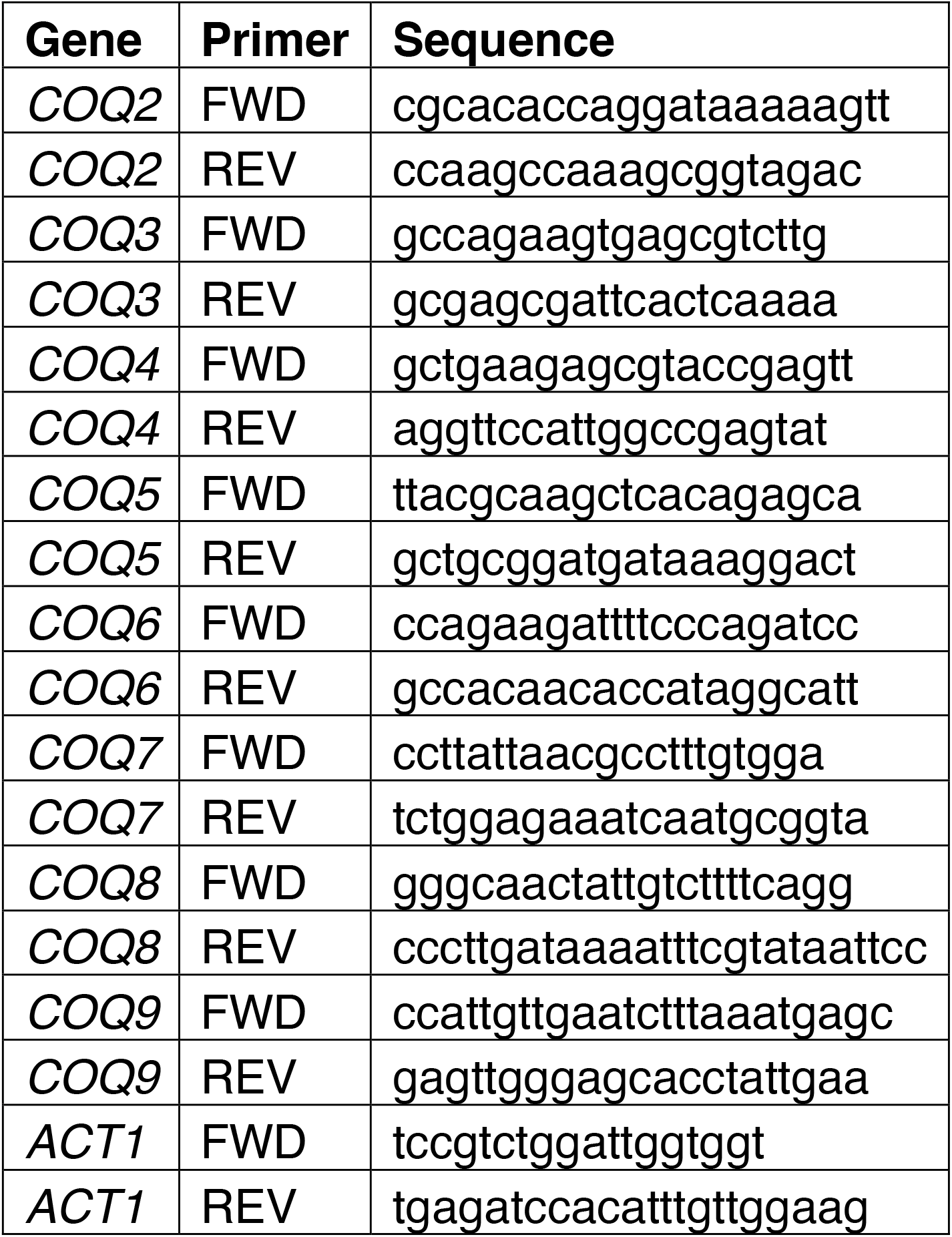
Primers for qPCR of *COQ* genes.

**Supplementary Table 2:**
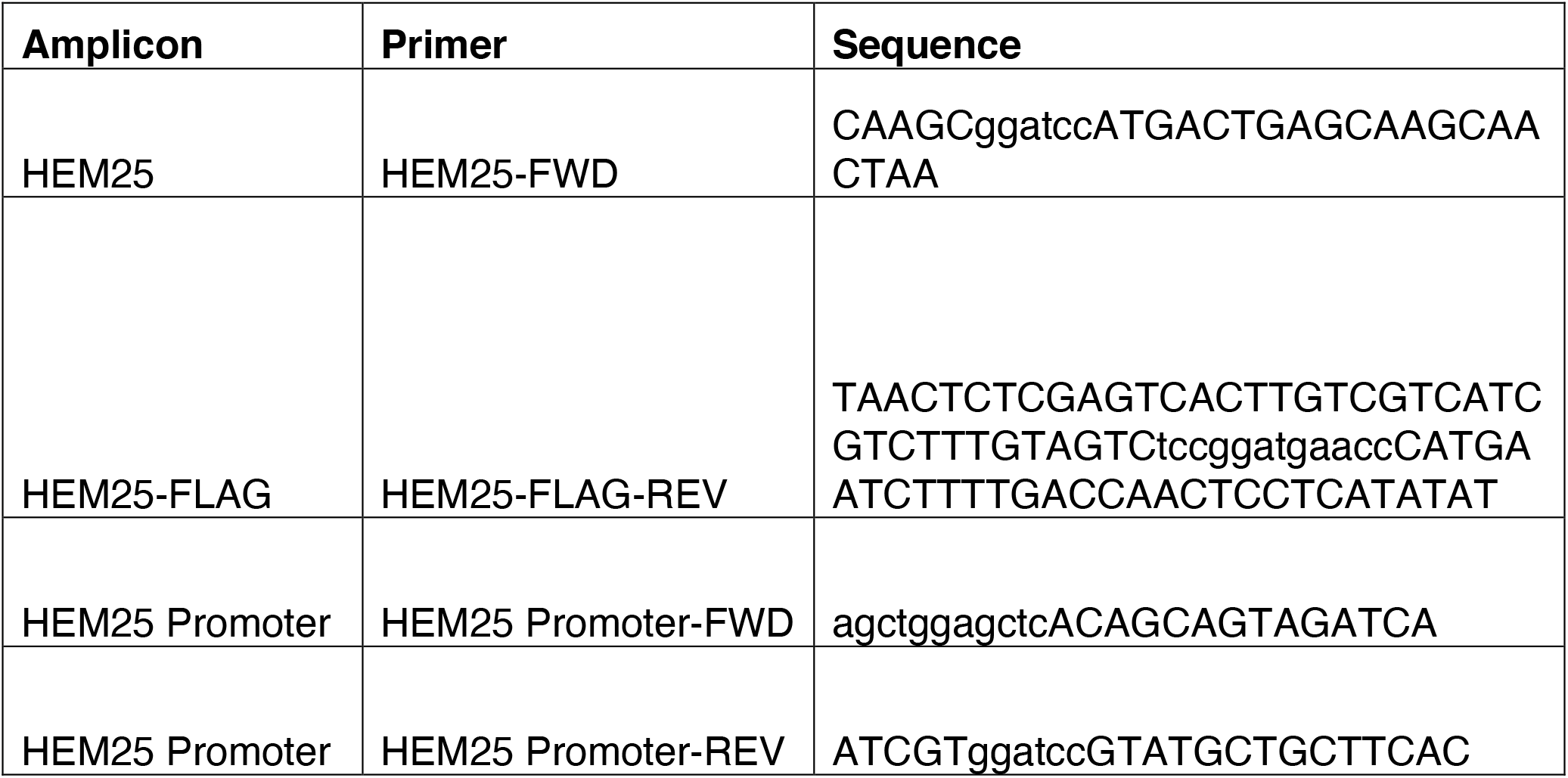
Primers used in this study for amplification.

**Supplementary Table 3:**
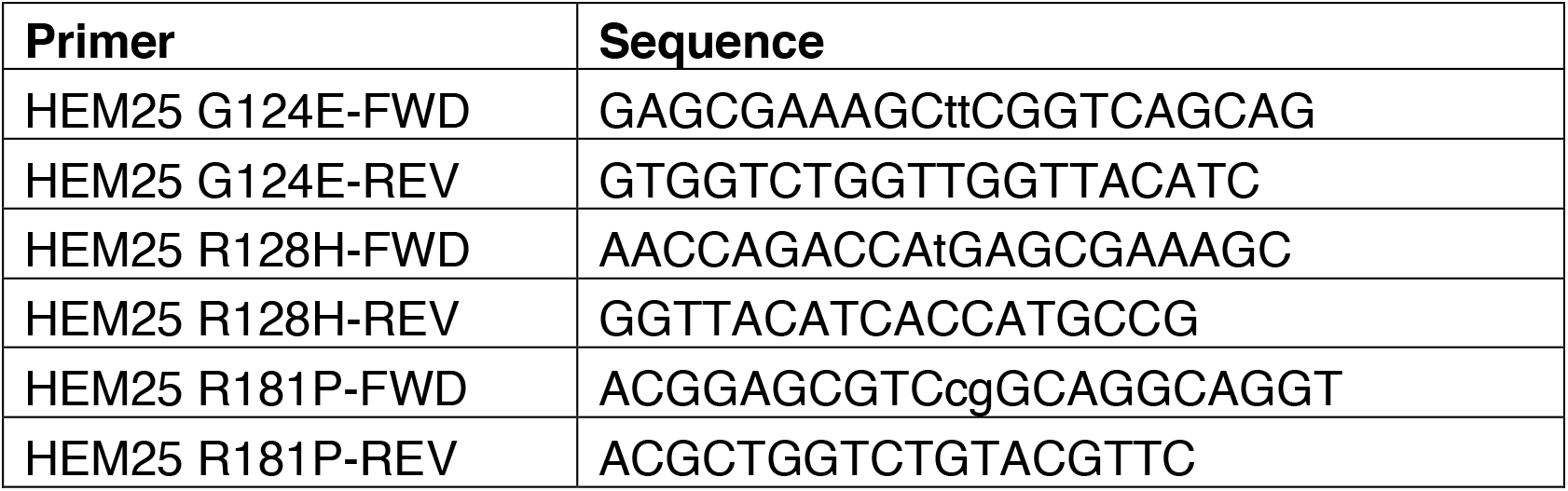
Primers used in this study for site directed mutagenesis.

## References

1. Lester, R. L. & Crane, F. L. The Natural Occurrence of Coenzyme Q and Related Compounds. f Biol Chem 234, 2169–2175 (1959).

2. Jones, M. E. Pyrimidine Nucleotide Biosynthesis in Animals: Genes, Enzymes, and Regulation of UMP Biosynthesis. Annu Rev Biochem 49, 253–279 (1980).

3. Frerman, F. E. Acyl-CoA dehydrogenases, electron transfer flavoprotein and electron transfer flavoprotein dehydrogenase. Biochem Soc T 16, 416–418 (1988).

4. Guerra, R. M. & Pagliarini, D. J. Coenzyme Q biochemistry and biosynthesis. Trends Biochem Sci (2023) doi:10.1016/j.tibs.2022.12.006.

5. Emmanuele, V. et al. Heterogeneity of Coenzyme Q10 Deficiency: Patient Study and Literature Review. Arch Neurol-chicago 69, 978–983 (2012).

6. Wang, Y. & Hekimi, S. The efficacy of coenzyme Q10 treatment in alleviating the symptoms of primary coenzyme Q10 deficiency: A systematic review. J Cell Mol Med 26, 4635–4644 (2022).

7. Stefely, J. A. & Pagliarini, D. J. Biochemistry of Mitochondrial Coenzyme Q Biosynthesis. Trends Biochem Sci 42, 824–843 (2017).

8. Awad, A. M. et al. Coenzyme Q10 deficiencies: pathways in yeast and humans. Essays Biochem 62, 361–376 (2018).

9. Wang, Y. & Hekimi, S. Understanding Ubiquinone. Trends Cell Biol 26, 367–378 (2016).

10. Gold, P. H. & Olson, R. E. Studies on Coenzyme Q. The biosynthesis of coenzyme Q9 in rat tissue slices. f Biol Chem 241, 3507–3516 (1966).

11. Payet, L. A. et al. Mechanistic Details of Early Steps in Coenzyme Q Biosynthesis Pathway in Yeast. Cell Chem Biol 23, 1241–1250 (2016).

12. Casey, J. & Threlfall, D. R. Synthesis of 5-demethoxyubiquinone-6 and ubiquinone-6 from 3-hexaprenyl-4-hydroxybenzoate in yeast mitochondria. Febs Lett 85, 249–253 (1978).

13. Momose, K. & Rudney, H. 3-Polyprenyl-4-hydroxybenzoate Synthesis in the Inner Membrane of Mitochondria from p-Hydroxybenzoate and Isopentenylpyrophosphate. A demonstration of isoprenoid synthesis in rat liver mitochondria. f Biol Chem 247, 3930–3940 (1972).

14. Tzagoloff, A. & Dieckmann, C. L. PET genes of Saccharomyces cerevisiae. Microbiol Rev 54, 211–225 (1990).

15. Robinson, K. P. et al. Defining intermediates and redundancies in coenzyme Q precursor biosynthesis. f Biological Chem 296, 100643 (2021).

16. Taylor, E. B. Functional Properties of the Mitochondrial Carrier System. Trends Cell Biol 27, 633–644 (2017).

17. Palmieri, F. & Monne, M. Discoveries, metabolic roles and diseases of mitochondrial carriers: A review. Biochimica Et Biophysica Acta Bba - Mol Cell Res 1863, 2362–78 (2016).

18. Vogtle, F. N. et al. Landscape of submitochondrial protein distribution. Nat Commun 8, 290 (2017).

19. Morgenstern, M. et al. Definition of a High-Confidence Mitochondrial Proteome at Quantitative Scale. Cell Reports 19, 2836–2852 (2017).

20. Ozeir, M. et al. Coenzyme Q biosynthesis: Coq6 is required for the C5-hydroxylation reaction and substrate analogs rescue Coq6 deficiency. Chem Biol 18, 1134–42 (2011).

21. Marobbio, C. M. T., Agrimi, G., Lasorsa, F. M. & Palmieri, F. Identification and functional reconstitution of yeast mitochondrial carrier for S-adenosylmethionine. Embo f 22, 5975–5982 (2003).

22. Barkovich, R. J. et al. Characterization of the COQ5 Gene from Saccharomyces cerevisiae EVIDENCE FOR A C-METHYLTRANSFERASE IN UBIQUINONE BIOSYNTHESIS^*^. f Biol Chem 272, 9182–9188 (1997).

23. Jonassen, T. & Clarke, C. F. Isolation and Functional Expression of Human COQ3, a Gene Encoding a Methyltransferase Required for Ubiquinone Biosynthesis^*^ J Biol Chem 275, 12381–12387 (2000).

24. Luk, E., Carroll, M., Baker, M. & Culotta, V. C. Manganese activation of superoxide dismutase 2 in Saccharomyces cerevisiae requires MTM1, a member of the mitochondrial carrier family. Proc National Acad Sci 100, 10353–10357 (2003).

25. Diessl, J. et al. Manganese-driven CoQ deficiency. Nat Commun 13, 6061 (2022).

26. Naranuntarat, A., Jensen, L. T., Pazicni, S., Penner-Hahn, J. E. & Culotta, V. C. The Interaction of Mitochondrial Iron with Manganese Superoxide Dismutase^*^. J Biol Chem 284, 22633–22640 (2009).

27. Lunetti, P. et al. Characterization of Human and Yeast Mitochondrial Glycine Carriers with Implications for Heme Biosynthesis and Anemia. J Biol Chem 291, 19746–59 (2016).

28. Fernandez-Murray, J. P. et al. Glycine and Folate Ameliorate Models of Congenital Sideroblastic Anemia. Plos Genet 12, e1005783 (2016).

29. Johnson, A. et al. COQ9, a New Gene Required for the Biosynthesis of Coenzyme Q in Saccharomyces cerevisiae ^*^. J Biol Chem 280, 31397–31404 (2005).

30. Stefely, J. A. et al. Mitochondrial protein functions elucidated by multi-omic mass spectrometry profiling. Nat Biotechnol 34, 1191–1197 (2016).

31. Lee, A. Y. et al. Mapping the Cellular Response to Small Molecules Using Chemogenomic Fitness Signatures. Science 344, 208–211 (2014).

32. Desbats, M. A. et al. The COQ2 genotype predicts the severity of coenzyme Q 10 deficiency. Hum Mol Genet 25, 4256–4265 (2016).

33. Allan, C. M. et al. Identification of Coq11, a New Coenzyme Q Biosynthetic Protein in the CoQ-Synthome in Saccharomyces cerevisiae^*^. J Biol Chem 290, 7517–7534 (2015).

34. Hsieh, E. J. et al. Saccharomyces cerevisiae Coq9 polypeptide is a subunit of the mitochondrial coenzyme Q biosynthetic complex. Arch Biochem Biophys 463, 19–26 (2007).

35. Dufay, J. N., Fernández-Murray, J. P. & McMaster, C. R. SLC25 Family Member Genetic Interactions Identify a Role for HEM25 in Yeast Electron Transport Chain Stability. G3 Genes Genomes Genetics 7, 1861–1873 (2017).

36. Ravaud, S. et al. Impaired Transport of Nucleotides in a Mitochondrial Carrier Explains Severe Human Genetic Diseases. Acs Chem Biol 7, 1164–1169 (2012).

37. Mifsud, J. et al. The substrate specificity of the human ADP/ATP carrier AAC1. Mol Membr Biol 30, 160–168 (2012).

38. Castegna, A. et al. Identification and Functional Characterization of a Novel Mitochondrial Carrier for Citrate and Oxoglutarate in Saccharomyces cerevisiae ^*^. J Biol Chem 285, 17359–17370 (2010).

39. Nichols, N. N. & Harwood, C. S. PcaK, a high-affinity permease for the aromatic compounds 4-hydroxybenzoate and protocatechuate from Pseudomonas putida. J Bacteriol 179, 5056–5061 (1997).

40. Robinson, A. J., Overy, C. & Kunji, E. R. The mechanism of transport by mitochondrial carriers based on analysis of symmetry. Proc National Acad Sci 105, 17766–71 (2008).

41. Guernsey, D. L. et al. Mutations in mitochondrial carrier family gene SLC25A38 cause nonsyndromic autosomal recessive congenital sideroblastic anemia. Nat Genet 41, 651–3 (2009).

42. Heeney, M. M. et al. SLC25A38 congenital sideroblastic anemia: Phenotypes and genotypes of 31 individuals from 24 families, including 11 novel mutations, and a review of the literature. Hum. Mutat. 42, 1367–1383 (2021).

43. Rensvold, J. W. et al. Defining mitochondrial protein functions through deep multiomic profiling. Nature 606, 382–388 (2022).

44. Thomas, P. D. et al. PANTHER: Making genome-scale phylogenetics accessible to all. Protein Sci 31, 8–22 (2022).

45. Subramanian, K. et al. Coenzyme Q biosynthetic proteins assemble in a substrate-dependent manner into domains at ER-mitochondria contacts. J Cell Biol 218, 1353–1369 (2019).

46. Eisenberg-Bord, M. et al. The Endoplasmic Reticulum-Mitochondria Encounter Structure Complex Coordinates Coenzyme Q Biosynthesis. Contact 2, 2515256418825409 (2019).

47. Vest, K. E., Leary, S. C., Winge, D. R. & Cobine, P. A. Copper Import into the Mitochondrial Matrix in Saccharomyces cerevisiae Is Mediated by Pic2, a Mitochondrial Carrier Family Protein*. J Biol Chem 288, 23884–23892 (2013).

48. Boulet, A. et al. The mammalian phosphate carrier SLC25A3 is a mitochondrial copper transporter required for cytochrome c oxidase biogenesis. J Biol Chem 293, 1887–1896 (2018).

49. Casey, J. & Threlfall, D. R. Formation of 3-hexaprenyl-4-hydroxybenzoate by matrix-free mitochondrial membrane-rich preparations of yeast. Biochim Biophys Acta 530, 487–502 (1978).

50. Runquist, M., Ericsson, J., Thelin, A., Chojnacki, T. & Dallner, G. Isoprenoid biosynthesis in rat liver mitochondria. Studies on farnesyl pyrophosphate synthase and trans-prenyltransferase.J Biol Chem 269, 5804–5809 (1994).

51. Grünler, J., Ericsson, J. & Dallner, G. Branch-point reactions in the biosynthesis of cholesterol, dolichol, ubiquinone and prenylated proteins. Biochimica Et Biophysica Acta Bba - Lipids Lipid Metabolism 1212, 259–277 (1994).

52. Keller, R. K. Squalene synthase inhibition alters metabolism of nonsterols in rat liver. Biochimica Et Biophysica Acta Bba - Lipids Lipid Metabolism 1303, 169–179 (1996).

53. Kardon, J. R. et al. Mitochondrial ClpX Activates a Key Enzyme for Heme Biosynthesis and Erythropoiesis. Cell 161, 858–867 (2015).

54. Rondelli, C. M. et al. The ubiquitous mitochondrial protein unfoldase CLPX regulates erythroid heme synthesis by control of iron utilization and heme synthesis enzyme activation and turnover. J Biol Chem 297, 100972 (2021).

55. Stefely, J. A. et al. Cerebellar Ataxia and Coenzyme Q Deficiency through Loss of Unorthodox Kinase Activity. Mol Cell 63, 608–620 (2016).

56. Ziegler, M. et al. Welcome to the Family: Identification of the NAD+ Transporter of Animal Mitochondria as Member of the Solute Carrier Family SLC25. Biomol 11, 880 (2021).

57. Luongo, T. S. et al. SLC25A51 is a mammalian mitochondrial NAD+ transporter. Nature 588, 174–179 (2020).

58. Kory, N. et al. MCART1/SLC25A51 is required for mitochondrial NAD transport. Sci Adv 6, eabe5310 (2020).

59. Girardi, E. et al. Epistasis-driven identification of SLC25A51 as a regulator of human mitochondrial NAD import. Nat Commun 11, 6145 (2020).

## Methods references

60. Baudin, A., Ozier-Kalogeropoulos, O., Denouel, A., Lacroute, F. & Cullin, C. A simple and efficient method for direct gene deletion in Saccharomyces cerevisiae. Nucleic Acids Res 21, 3329–3330 (1993).

61. Longtine, M. S. et al. Additional modules for versatile and economical PCR-based gene deletion and modification in Saccharomyces cerevisiae. Yeast 14, 953–961 (1998).

62. Wach, A., Brachat, A., Alberti-Segui, C., Rebischung, C. & Philippsen, P. Heterologous HIS3 Marker and GFP Reporter Modules for PCR-Targeting in Saccharomyces cerevisiae. Yeast 13, 1065–1075 (1997).

63. Ismail, A. et al. Coenzyme Q Biosynthesis: Evidence for a Substrate Access Channel in the FAD-Dependent Monooxygenase Coq6. Plos Comput Biol 12, e1004690 (2016).

64. Sassa, S. Sequential induction of heme pathway enzymes during erythroid differentiation of mouse Friend leukemia virus-infected cells. J Exp Medicine 143, 305–315 (1976).

65. Sinclair, P. R., Gorman, N. & Jacobs, J. M. Measurement of Heme Concentration. Curr Protoc Toxicol 00, 8.3.1-8.3.7 (1999).

66. Shishkova, E., Hebert, A. S., Westphall, M. S. & Coon, J. J. Ultra-High Pressure (>30,000 psi) Packing of Capillary Columns Enhancing Depth of Shotgun Proteomic Analyses. Anal Chem 90, 11503–11508 (2018).

67. Hebert, A. S. et al. Improved Precursor Characterization for Data-Dependent Mass Spectrometry. Anal Chem 90, 2333–2340 (2018).

68. Tyanova, S., Temu, T. & Cox, J. The MaxQuant computational platform for mass spectrometry-based shotgun proteomics. Nat Protoc 11, 2301–2319 (2016).

69. Cox, J. et al. Accurate Proteome-wide Label-free Quantification by Delayed Normalization and Maximal Peptide Ratio Extraction, Termed MaxLFQ*. Mol Cell Proteomics 13, 2513–2526 (2014).

70. Brademan, D. R. et al. Argonaut: A Web Platform for Collaborative Multi-omic Data Visualization and Exploration. Patterns 1, 100122 (2020).

71. Meisinger, C., Pfanner, N. & Truscott, K. N. Isolation of yeast mitochondria. Methods Mol Biol 313, 33–9 (2006).

72. Miroux, B. & Walker, J. E. Over-production of Proteins inEscherichia coli: Mutant Hosts that Allow Synthesis of some Membrane Proteins and Globular Proteins at High Levels. J Mol Biol 260, 289–298 (1996).

73. Gietz, R. D. & Schiestl, R. H. High-efficiency yeast transformation using the LiAc/SS carrier DNA/PEG method. Nat Protoc 2, 31–34 (2007).

74. Waterhouse, A. M., Procter, J. B., Martin, D. M. A., Clamp, M. & Barton, G. J. Jalview Version 2—a multiple sequence alignment editor and analysis workbench. Bioinformatics 25, 1189–1191 (2009).

75. Sievers, F. et al. Fast, scalable generation of high-quality protein multiple sequence alignments using Clustal Omega. Mol Syst Biol 7, 539–539 (2011).

76. Varadi, M. et al. AlphaFold Protein Structure Database: massively expanding the structural coverage of protein-sequence space with high-accuracy models. Nucleic Acids Res 50, D439–D444 (2021).

77. Jumper, J. et al. Highly accurate protein structure prediction with AlphaFold. Nature 596, 583–589 (2021).

78. Veling, M. T. et al. Multi-omic Mitoprotease Profiling Defines a Role for Oct1p in Coenzyme Q Production. Mol Cell 68, 970-977.e11 (2017).

